# Chimeric MerR-Family Regulators and Logic Elements for the Design of Metal Sensitive Genetic Circuits in *Bacillus subtilis*

**DOI:** 10.1101/2022.10.13.512145

**Authors:** Jasdeep S. Ghataora, Susanne Gebhard, Bianca J. Reeksting

## Abstract

Whole-cell biosensors are emerging as promising tools for monitoring environmental pollutants such as heavy metals. These sensors constitute a genetic circuit comprising a sensing module and an output module, such that a detectable signal is produced in the presence of the desired analyte. The MerR family of metal-responsive regulators offers great potential for the construction of metal sensing circuits, due to their high sensitivity, tight transcription control and large diversity in metal-specificity. However, the sensing diversity is broadest in Gram-negative systems, while chassis organisms are often selected from Gram-positive species, particularly sporulating bacilli. This can be problematic, because Gram-negative biological parts, such as promoters, are frequently observed to be non-functional in Gram-positive hosts. Herein, we combined construction of synthetic genetic circuits and chimeric MerR regulators, supported by structure-guided design, to generate metal-sensitive biosensor modules that are functional in the biotechnological work-horse species *Bacillus subtilis*. These chimeras consist of a constant Gram-positive derived DNA-binding domain fused to variable metal binding domains of Gram-negative origins. To improve the specificity of the whole-cell biosensor, we developed a modular ‘AND gate’ logic system based on the *B. subtilis* natively split σ-factor, SigO-RsoA, designed to maximise future use for synthetic biology applications in *B. subtilis*. This work provides insights into the use of modular regulators, such as the MerR family, in the design of synthetic circuits for the detection of heavy metals, with potential wider applicability of the approach to other systems and genetic backgrounds.

## INTRODUCTION

Heavy metal pollution, caused by anthropogenic activities such as metallurgic processes associated with increased industrialisation and the overuse of pesticides and fertilisers, poses a risk to the environment and human health ^1,2^. These metals cannot be broken down and subsequently accumulate within the environment. Furthermore, the presence of heavy metals has been linked to the co-selection of antibiotic resistance genes, as resistance determinants for heavy metals and antibiotics frequently co-occur on mobile genetic elements ^3–5^. As a result, the persistence of such contaminants in waterways is likely to encourage the dissemination of antibiotic resistance genes in the environment ^6,7^. It is therefore important to monitor environmental levels of metal contaminants to identify and manage risks, as well as to implement and assess remediation strategies. Traditional analytical techniques such as Atomic Absorption Spectroscopy and Inductively Coupled Plasma Mass Spectrometry (ICP-MS) offer high sensitivity in detection of toxic metals in contaminated environments, but are hampered by cost, lack of *in situ* monitoring, and the requirement for trained personnel. They are also unable to specifically report the biologically available fractions of polluting metals, which present the most direct risk to human or environmental health. Advances in synthetic biology in combination with decreasing costs of DNA synthesis have made whole-cell biosensors, based on a microbial chassis into which a genetic circuit is built for the detection of an analyte of interest, a viable option to circumvent these limitations. Indeed, the construction of synthetic circuits in bacteria to develop whole cell biosensors for the monitoring of heavy metals has gained considerable interest ^8–11^.

Metalloregulatory systems offer a source of biological parts for the construction of whole-cell biosensors sensitive to heavy metals ^12–14^. The MerR protein family is a well described example of metal-responsive regulators ^15,16^. The corresponding target promoters are characterised by an unusually long spacer region (19-20 bp) between the -10 and -35 elements of a σ^70^/σ^A^ dependent promoter, which places these elements on opposite faces of the DNA. As a result, the promoters are a poor substrate for RNA polymerase binding and transcription initiation ^17^. The regulator, MerR, binds between the -10 and -35 elements of the promoter and upon binding of an inducer, such as Hg^2+^ or Cu^+^ ions, undergoes a conformational change to under-twist the promoter and re-align the - 10 and -35 elements. This facilitates recognition by RNA polymerase and triggers transcription initiation ^17,18^. The tightly controlled mechanism of transcription and the high sensitivity and selectivity of MerR regulators for specific metal ions make them ideal for the design of metal sensing circuits ^8,9,19^. Moreover, as MerR proteins are located cytoplasmically, toxic metals must pass into the cell to evoke a transcriptional response. MerR-based biosensors thus give an indication of the bioavailability of a given contaminant.

MerR regulators have a modular architecture consisting of two discrete domains, an N-terminal DNA binding domain (DBD) responsible for promoter recognition ^20–22^ and a C-terminal domain with a metal binding loop for the coordination of metal ions, referred to here as the metal binding domain (MBD) ^23–25^. The specificity of metal recognition in the MBD is determined by metal coordinating amino acids that allow the coordination of some metals but exclude others. Diversity within these domains facilitates the detection of different metals, providing potential candidates for biosensors with different specificity. These MerR proteins can be used as the sensory modules in biosensors, with their corresponding target promoter fused to a detectible output, e.g. fluorescent or luminescent reporter genes. However, harnessing the sensing diversity of the MerR family requires incorporating a new protein every time the specificity needs to be changed, each of which includes a new DBD that recognises a different promoter. This necessitates the redesign of the output module to ensure the promoter is recognized, and a signal can be detected.

Furthermore, the largest diversity of metal-specificity in MerR family regulators is found in Gram-negative bacteria –including ZntR (for Zn^2+^) and CueR (for Cu^+^) ^26^. Heterologous use of regulators in chassis systems from unrelated species can be problematic due to competition for host transcription and translation machinery resources ^27,28^, interference from the host genetic background ^29^, and species-specific differences in the recognition of circuit parts, such as promoters, ribosome binding sites and other regulatory features ^30–32^. Synthetic biology approaches can circumvent these problems, for example through rewiring biological circuits with synthetic promoters to solve transcriptional incompatibilities. It was also shown that a chimeric two-component response regulator produced by domain-swapping could restore functionality, such as seen with *Escherichia coli* derived NarX-NarL for use in *Bacillus subtilis* ^33^. The modular nature of MerR-family regulators may make these proteins well-suited to such approaches.

The Gram-positive, spore-forming bacterium *Bacillus subtilis* represents an ideal candidate as a chassis for whole cell biosensors given its biotechnological relevance, genetic tractability and availability of extensive genetic resources ^34–36^. Numerous examples exist of synthetic biosensor circuits that have been implemented in this chassis organism with application in the detection of pathological biomarkers, antibiotics, antifungal polyenes, and parasites ^37–40^. Based on these features, we here aimed to use *B. subtilis* as the host organism to explore the possibility of using domain swapping to engineer MerR-based biosensor circuits with a range of specificities. We hypothesised that combining variable MBDs with a constant DBD that is functional in *B. subtilis* would allow us to harness the metal binding diversity of proteins from Gram-negative, while ensuring compatibility with the hosts’ transcriptional machinery. Biosensor design would further be simplified using a single, luciferase-based output module.

An additional challenge in developing application-relevant biosensors is the potentially broad substrate specificity of some MerR regulators, which respond to multiple heavy metals and thus do not facilitate differentiation between specific contaminants. A solution for this may be found in logic gates, such as AND gates, which can be introduced into synthetic circuits to improve specificity ^9^. AND gates require multiple separate inputs to produce an output. In this way, combinations of non-specific sensing modules can be assembled to produce specific sensors. The use of recombinase-based AND logic circuits has been demonstrated, but these suffer from slow response times, which limits application ^37^. Whilst many examples of AND gates in the Gram-negative bacterium *E. coli* exist^29,41,42^, there are comparatively fewer examples in *B. subtilis*. Based on the extensive genetic resources available for *B. subtilis*, we here sought to design a logic gate to circumvent slow response times as well as improve the specificity of our designed sensors.

The present study describes the design, optimisation, and characterisation of heavy metal biosensors in *B. subtilis* based on chimeric MerR transcription factors. We first demonstrate the functionality of a heterologous MerR circuit derived from *Staphylococcus aureus* TW20 ^43^ in *B. subtilis* for the detection of Hg^2+^ ions. Subsequently, domain-swapping with two representative Gram-negative MerR-family regulators, ZntR (*E*. coli) and CueR (*E. coli)*, is used to engineer novel specificity of metal detection in *B. subtilis*. We demonstrate that rational engineering, guided by protein structure predictions, can be used to improve the functionality of such hybrids. To overcome problems with cross-specificity we engineer a standardised and modular AND gate logic system based on the *B. subtilis* natively split σ-factor system, SigO-RsoA – demonstrating its use in generating an ultra-specific heavy metal detection circuit. Our results establish the basic design rules for functional hybrid MerR-based metal sensors, which should easily be adaptable to broaden the range of detectable metals in the *B. subtilis* chassis and may enable construction of functional hybrid regulators in other genetic backgrounds.

## RESULTS AND DISCUSSION

### Functional reconstitution of a heterologous Hg^2+^-sensitive circuit in *B. subtilis* W168

*Bacillus subtilis* lacks a native metal-sensitive MerR regulatory system. We thus first sought to construct a synthetic metal-sensitive circuit using the regulatory components of the MerR mercury resistance determinant from *S. aureus* TW20 ^43^. This included the regulator MerR and its cognate promoter P_*merR20*_, which we combined into the whole-cell biosensor genetic circuit shown in Figure 1A. In this circuit, in the absence of any inducing Hg^2+^ ions, MerR should bind and repress the P_*merR20*_ promoter which possesses an elongated spacer region between the -10 and -35 elements, rendering it a poor substrate for RNAP recognition. As P_*merR20*_ was fused to the bacterial luciferase operon *luxABCDE*, no luminescence should be detected under these conditions. Binding of Hg^2+^ (input) promotes conformational changes within MerR ^44^ (whose production is driven by the xylose-inducible promoter, P_*xylA*_), facilitating under-twisting of the promoter (P_*merR20*_) and re-aligning the -10 and -35 element, allowing for RNAP to initiate expression of *luxABCDE* (output). Thus, luciferase activity of the cells should be correlated with the concentration of exogenous Hg^2+^ ions.

**Figure 1.**
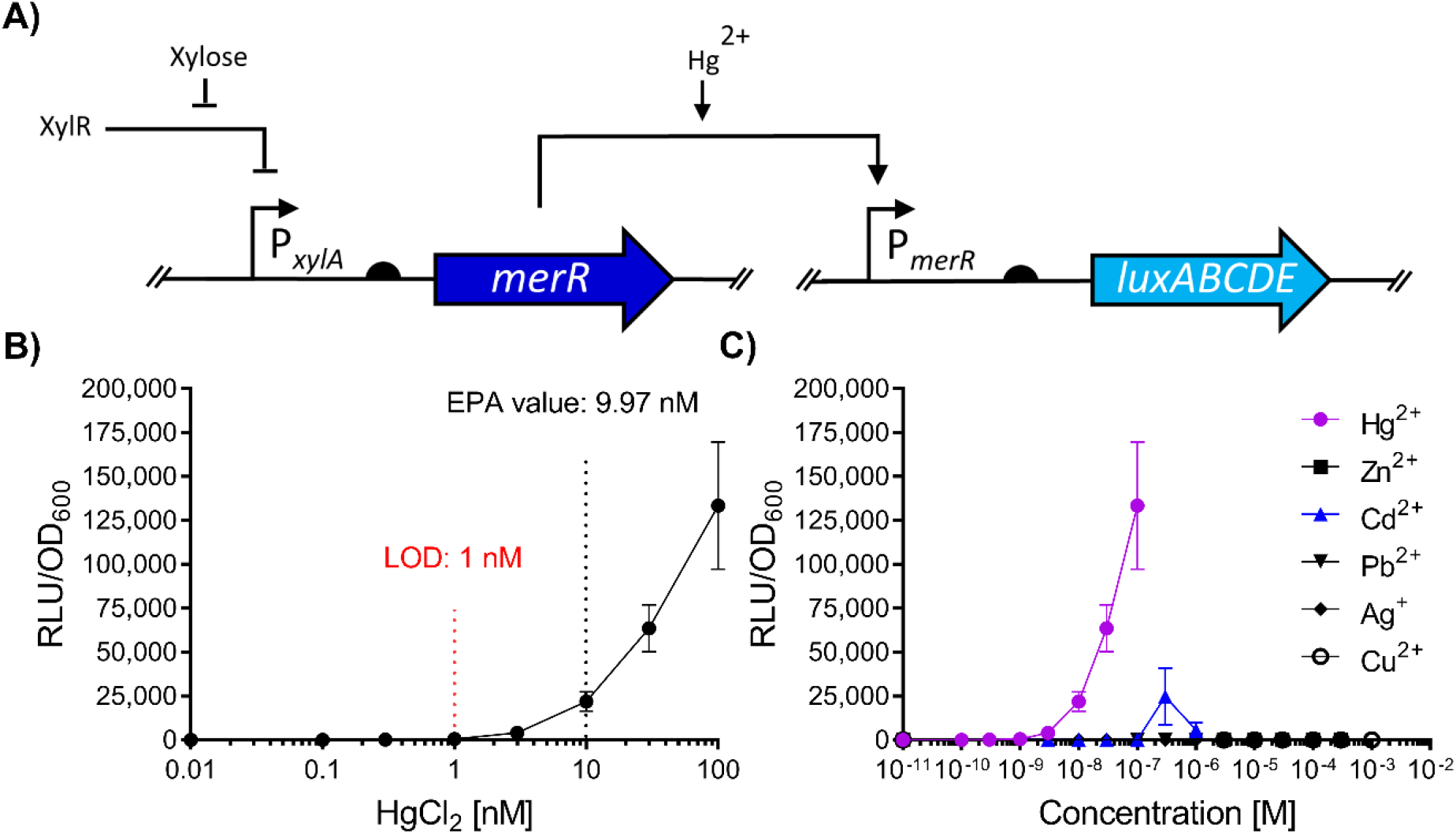
Design and evaluation of a heterologous Hg^2+^-sensing synthetic circuit in *B. subtilis*. Schematic representation of the MerR synthetic sensing module in *B. subtilis* W168 derived from *S. aureus* TW20. In the circuit, induction with Hg^2+^ allows MerR to activate luciferase expression as a function of Hg^2+^ concentration. Bent arrows indicate promoters, flat-head arrows indicate inhibition, black semi-circles indicate ribosome binding sites and genes for both *merR* and *luxABCDE* coding sequences are indicated by dark and light blue arrows, respectively. **B)** Dose response of the Hg^2+^ circuit. The Environmental protection agency (EPA) guideline value is indicated, and the estimated limit of detection (LOD) for the regulatory circuit is shown (red). **C)** Metal specificity of the circuit. For panels **(B-C)**, cells were grown to OD_600_ =∼0.03 and induced with the concentrations of metals as indicated with luciferase activity output (relative luminescence units [RLU]) normalised to optical density (OD_600_) values (RLU/OD_600_) for three time points (35, 40 and 45 mins) post-induction. Values are presented as mean and ± standard deviation of either two or three independent replicates.

To test the functionality of this circuit, *B. subtilis* cells harbouring both the P_*xylA*_-*merR* and the P_*merR20*_-*luxABCDE* constructs (SGB1005) were challenged with sub-inhibitory concentrations of Hg^2+^, and promoter activity was monitored. The resulting dose-response behaviour (Fig. 1B) showed that promoter activity (RLU/OD_600_) was proportional to the Hg^2+^concentration with a limit of detection (LOD) of 1 nM, below the guideline values set by the Environmental Protection Agency (9.97 nM). The sensitivity of this first, simple biosensor was already comparable to previous Hg^2+^-inducible systems, which had required RBS tuning and amplification circuits to achieve such sensitivity ^8^. Therefore, we have demonstrated that a heterologous MerR resistance determinant from the Gram-positive *S. aureus* was functional as part of a synthetic circuit in *B. subtilis*. In addition, the operational range of this circuit was relevant to application and within the range of values (0.99 - 17.95 nM) found in environmental samples according to global surveys of total Hg^2+^ in water over the last 50 years ^45^.

To determine the metal specificity of the circuit we first determined the MIC values of a range of heavy metals (MICs: Cd^2+^ [10 μM], Ag^+^ [10 μM], Zn^2+^ [1 mM], Cu^+^ [3 mM], Pb^2+^ [100 μM] and Hg^2+^ [1000 nM]) and subsequently tested our circuit using sub-lethal concentrations of these metals (Fig. 1C). Strong induction (35-45 mins after metal addition) was seen for Hg^2+^, whereas Zn^2+^, Cu^2+^, and Pb^2+^ did not produce significant signals. The addition of cadmium (Cd^2+^) at 0.3 μM led to a detectable signal and at higher levels of Cd^2+^ (1 μM), a drop in RLU/OD_600_ activity was observed (Fig. 1C). Heavy metals such as Cd^2+^ are known to disrupt protein folding, which includes proteins such as luciferases^46^. Therefore, these results could indicate inhibitory effects of Cd^2+^ on reporter output at this concentration, despite being ten-fold below MIC. As Cd^2+^ concentrations in the environment over the last 20 years cover a range from ∼ 0.02 – 5 μM ^47^, these may realistically be detected by our sensory circuit. We therefore considered the response profile for this sensing module to be specific for Hg^2+^ with some cross-reactivity to Cd^2+^.

### Guided design of hybrid regulators to alter metal-specificity of the sensing module

Having established a functional sensing module, we then wanted to further explore the use of MerR homologues to construct sensors in *B. subtilis* with specificities for other metals. Phylogenetic analysis shows that in Gram-positive bacteria, the specificity of MerR regulators appears to be restricted to Hg^2+^, whereas the diversity of metal specificity appears to be much broader in MerR regulators from Gram-negative species, including metals such as Zn^2+^, Cu^+^, Pb^2+^ and Cd^2+ 26^. Upon initial construction of metal detection circuits based on Gram-negative-derived biological parts in *B. subtilis*, we found these promoters to be non-functional in the *B. subtilis* host (Supplementary Fig. S1). This is consistent with several other studies that tested promoters from Gram-negatives in *B. subtilis*, including P_*tac*_ ^48^, P_*lacUV5*_ ^49^, the strong synthetic Anderson promoter J23101 ^31^, and the NarX-NarL two-component system target promoter P_*dcuS77*_ ^33^. This is likely due to differences in the transcription machinery, for example in σ-factor stringency between Gram-negative and Gram-positive species ^50^.

To circumvent the issue of host/biological part incompatibility, we investigated the possibility of exploiting the modularity of MerR regulators by using a domain-swap strategy to engineer the metal specificity of the sensing module. We had already demonstrated the compatibility of the *S. aureus* MerR protein and its target promoter P_*merR20*_ with the *B. subtilis* host, where the MerR DNA-binding domain (MerR_DBD_) determined the promoter specificity (see above). We therefore speculated that replacement of the MerR C-terminal metal binding region (conferring metal-specificity) with the corresponding region derived from a MerR homologue of Gram-negative origin may enable us to change the specificity of the sensing module. Indeed, such approaches have been successful using other small one-component regulators such as TetR, LacI and GalR, for a variety of synthetic circuit applications ^51,52^, as well as resolving part incompatibility in *B. subtilis* using chimeric two-component response regulators ^33^.

To test this approach, we designed a chimeric MerR regulator, MerRZntR, with the DNA-binding domain derived from the Hg^2+^-responsive MerR of *S. aureus* TW20 used above (MerR_DBD_; residues Met1-Tyr38) and the C-terminal metal binding domain derived from the Zn^2+^-responsive MerR homologue ZntR from *E. coli* (ZntR_MBD_; residues Arg38-Cys141) (Fig. 2A). The junction point of Arg38 (ZntR) between the MerR_DBD_ and ZntR_DBD_ was selected based on previous work on ZntR, which showed this region was susceptible to cleavage by trypsin and could separate ZntR into two domains^23^. The resulting chimeric amino acid sequence was used to generate a predicted homology model for the structure of MerRZntR, the monomer of which is shown for simplicity (Fig. 2B). The analysis of the homology model indicated an overall topology for MerRZntR similar to other MerR regulators, suggesting that the fusion of the two domains would be unlikely to disrupt the overall protein architecture.

**Figure 2.**
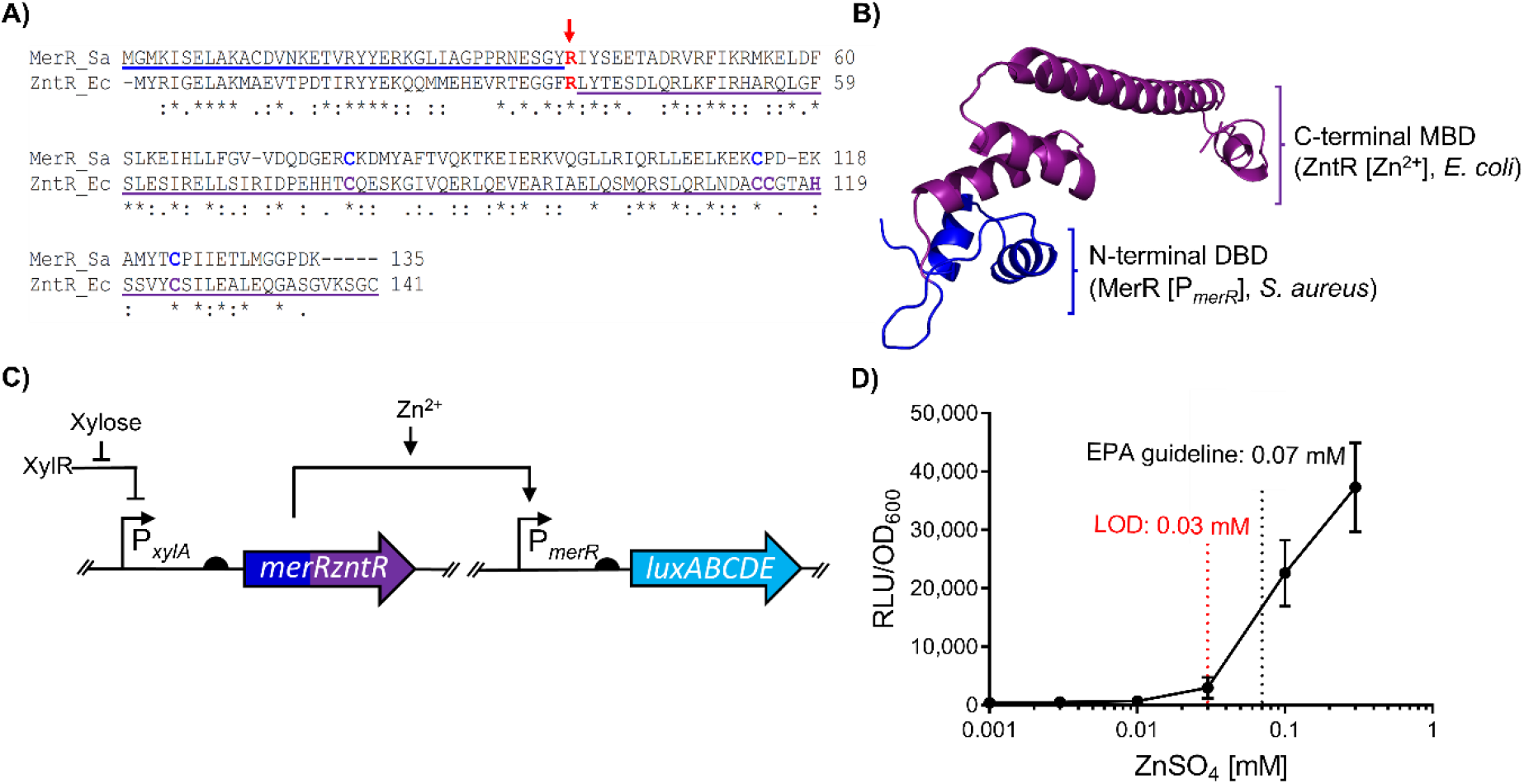
Design and assessment of the chimera, MerRZntR. **A)** Sequence alignment of MerR homologues. Both regulators MerR (*S. aureus*, “Sa”; accession code: CBI50741.1) and ZntR (*E. coli*, “Ec”; accession code: AAC76317.1) were aligned using the ClustalOmega tool^53^. The MerR derived DNA-Binding Domain (residues 1-38) is underlined blue, whilst the ZntR derived Metal-Binding Domain (residues 38 – 141) is underlined purple. The fusion point (Arg38, ZntR) is indicated in red with an arrow. Residues involve in Hg^2+^ coordination by MerR are indicated in dark blue, those involved in Zn^2+^ coordination by ZntR are indicated in purple. Asterisk “*” indicate fully conserved residues, colon “:” indicates conserved residues with similar properties, and period “.” indicate residues of weakly similar properties. **B)** The resulting homology model of MerRZntR using the aforementioned sequence in panel **(A)** generated using I-TASSER ^54^ is shown; the top-ranking structural analogue was CueR from *E. coli* (PDB:1Q05, C-score = 0.71, TM-score = 0.87). The origin of each domain, and their ligands are indicated using the same colour scheme as in A. **C)** Circuit schematic comprising the designed chimera MerRZntR. Bent arrows indicate promoters, flat-head arrows indicate inhibition, black semi-circles indicate ribosome binding sites, and genes for both *merRzntR* and *luxABCDE* coding sequences are indicated by the dark blue/purple and light blue arrows, respectively. **D)** Dose response of the MerRZntR chimera sensory circuit. Cells were grown to OD_600_ =∼0.03 and induced with the concentrations of Zn^2+^ indicated, with luciferase activity output (relative luminescence units [RLU]) normalised to optical density (OD_600_) values (RLU/OD_600_) recorded and averaged for three time points (35, 40 and 45 mins) post-induction. Values are presented as mean ± standard deviation of two or three independent replicates.

To test its functionality, the DNA sequence encoding the chimeric MerRZntR regulator was incorporated into the sensing module developed above, again under control of P_*xylA*_, and integrated into the *B. subtilis* chromosome. Activity of the chimeric protein was again monitored by its ability to control *luxABCDE* expression from P_*merR20*_ (Fig. 2C). The resulting strain (SGB1011) was then tested by measuring promoter activities of cells in exponential growth phase challenged with sub-lethal concentrations of Zn^2+^ (Fig. 2D). The results showed a Zn^2+^-concentration dependent response of the promoter, with an LOD of 0.03 mM and a response of 90-fold over unchallenged cells at 0.3 mM (Fig. 2D). This clearly indicated that a functional chimera was produced, with the domain-swap leading to a change in metal specificity so that the module could now sense Zn^2+^. The sensitivity of the module was relevant to environmental levels of contamination, with the EPA guideline value of 0.07 mM falling within the sensitivity range. The data thus demonstrate the feasibility of engineering novel heterologous metal-sensitive biological parts using domain swaps to introduce novel metal specificity into MerR type regulators and overcome problems such as promoter incompatibility in the host organism.

### Structure-guided mutagenesis allows optimisation of MerRZntR activity

Inter-domain amino acid communication within metal-sensitive transcription factors plays a key role in coordinating the binding of a metal with changes in DNA-binding to either activate or repress transcription ^55^. Examples of such communication can be seen for MerR as well as MerR homologues CueR and SoxR (*E. coli*), where formation of a hydrogen bonding network upon ligand binding mediates communication between the MBD and DBD to activate transcription by remodelling local promoter topology ^17,56^. To assess whether this type of inter-domain interaction was likely to have been affected by the construction of the MerRZntR hybrid, a homology model of native ZntR was generated (Fig. 3A). This was necessary, because the available partial structure of the protein (PDB:1Q08) lacked both the ZntR_DBD_ and inter-domain interactions with its MBD. Hydrogen bonding was observed in the homology model between residues Ser43_MBD_, Arg47_MBD_ and Glu28_DBD_ (Fig. 3A). However, in the chimera MerRZntR, Ala29_DBD_ is present at the position equivalent to the negatively charged Glu28 in ZntR (Supplementary Fig. S2A-B). Seeing as the interaction between Ser43_MBD_ and Glu28_DBD_ may potentially contribute to the signalling mechanism between the two domains, we created an Ala29Glu variant of the MerRZntR hybrid to determine whether we could improve the Zn^2+^ response. The variant MerRZntR^A29E^ (in strain SGB1035) indeed exhibited greater increases in luminescence across all tested concentrations when compared to MerRZntR, with an improved LOD of 0.01 mM Zn^2+^ compared to 0.03 mM and a maximum response of 128-fold at 0.3 mM (Fig. 3B) compared to 90-fold in MerRZntR. Therefore, restoring the hydrogen bonding between the two protein domains could indeed improve the protein’s activity. Interestingly, upon introduction of additional substitutions in the chimera MerRZntR to re-establish structural interactions, the variants MerRZntR^A29E/G30H^ and MerRZntR^A29E/G30H/P32V^ showed no discernible difference in RLU/OD_600_ when compared to the single variant A29E (Supplementary Fig. S3).

**Figure 3.**
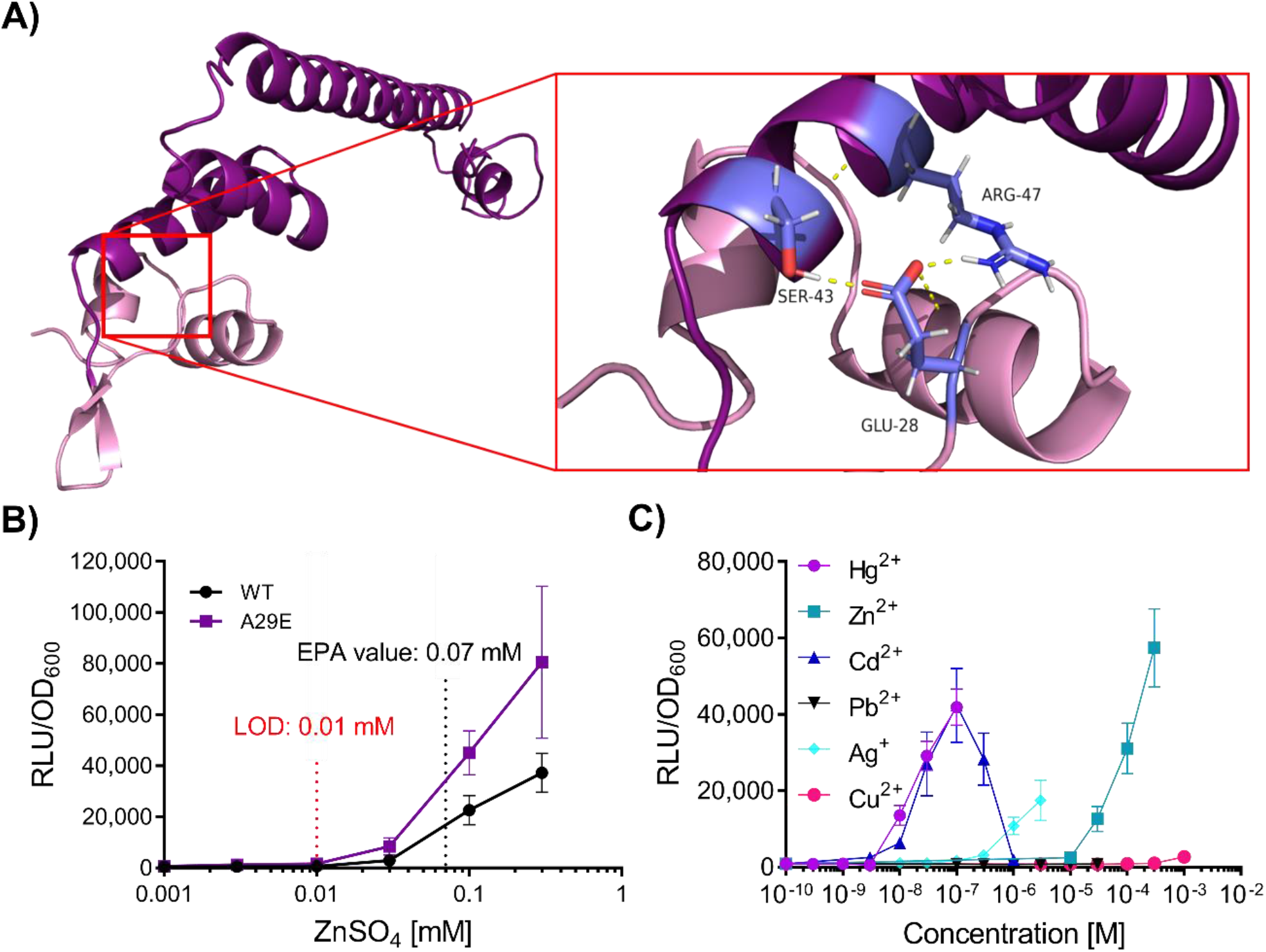
Structural analysis of ZntR to guide design of the chimera MerRZntR^A29E^. **A)** Homology model of ZntR. I-TASSER ^54^ was used to generate a full-length homology model of ZntR. *E. coli*, accession code : AAC76317.1, C-score = 0.71, TM score = 0.91). The DNA-Binding Domain is indicated in mauve, whilst the C-terminal Domain comprising the metal binding loop is indicated in dark purple. Residues involved in inter-domain communication (Glu-28, DNA-Binding Domain; Ser-43 and Arg-47, Metal-Binding Domain) are shown in lavender blue with hydrogen bonds shown in yellow. **B)** Guided mutagenesis of MerRZntR α-helix 2 and α-helix 3 loop to generate mutant MerRZntR^A29E^. The Environmental protection agency (EPA) guideline value is indicated, and the estimated limit of detection (LOD) for the regulatory circuit is shown (red). The most potent activator, MerRZntR^A29E^ is indicated in purple. **C)** Metal-specificity of the MerRZntR^A29E^ based circuit. Inducers are coloured using the key shown. For panels **(B-C)**, cells were grown to OD_600_ =∼0.03 and induced with the concentrations of metals as indicated, with luciferase activity output (relative luminescence units [RLU]) normalised to optical density (OD_600_) values (RLU/OD_600_) for three time points (35, 40 and 45 mins) post-induction. Values are presented as mean and ± standard deviation of either two or three independent replicates.

After selecting MerRZntR^A29E^ as the improved Zn^2+^ sensor, we sought to determine its substrate specificity profile. Strain SGB1035 was subjected to sub-inhibitory concentrations of various heavy metals (Fig. 3C). Cross-specificity was observed for MerRZntR^A29E^ to various heavy metals, including Hg^2+^, Cd^2+^, and to a lesser degree Ag^+^. This cross-specificity could be problematic in environments where these metals are likely to be co-contaminants, as it would make it impossible to distinguish between them and to estimate relevant levels of pollution. Taken together, our data indicate that structure-guided design is a promising approach for optimisation of newly designed chimeric biological parts – an approach we have utilised to improve both the overall RLU/OD_600_ output and LOD of a functional heterologous Zn^2+^ inducible circuit. However, the cross-specificity of the hybrid MerRZntR_A29E_ has downstream implications for specific monitoring of target heavy metals in potential field applications, which is further addressed below.

### Design and optimisation of a copper responsive hybrid MerRCueR through structure-guided mutagenesis

Having demonstrated the feasibility in principle of the domain-swap strategy, we investigated whether this approach could be applied to additional MerR homologues. To test this, we selected the well-characterized copper-responsive MerR homologue CueR of *E. coli* ^17^. A chimeric MerRCueR regulator was designed using amino acids 1-38 of MerR (*S. aureus*), as above, and residues 37-135 of CueR (*E. coli*), with Arg37 of CueR used as a fusion point between the two domains (Fig. 4A). The resulting chimeric amino acid sequence was used to generate a homology model of MerRCueR as before, with a C-score of 0.71 and TM-score of 0.81 (Fig. 4B), indicating a close match to the structure of CueR (PDB:1Q05). The circuit was then assembled in *B. subtilis* as above, using the P_*merR20*_-*luxABCDE* reporter (in strain SGB1027) to test functionality (Fig. 4C).

**Figure 4.**
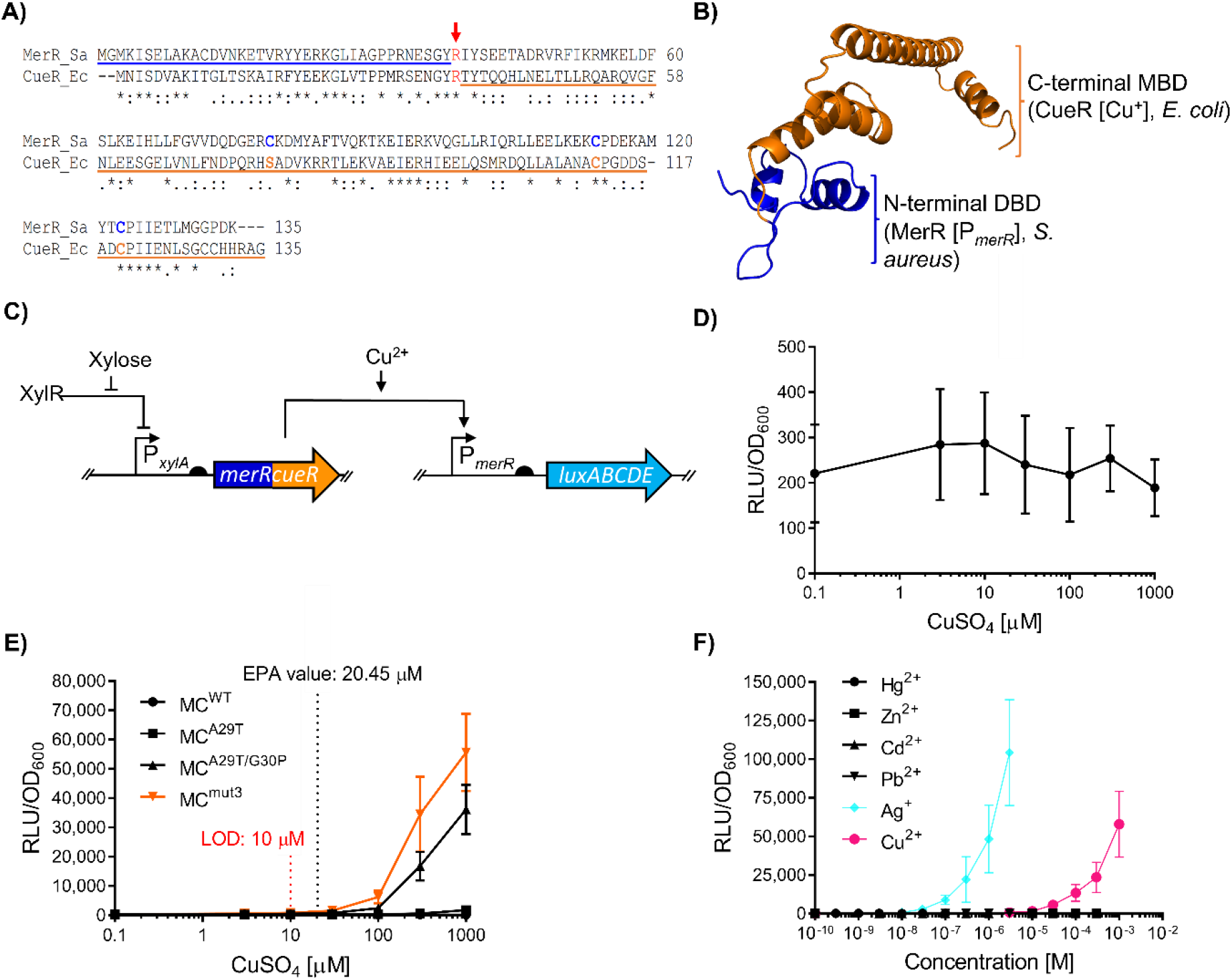
Design, assessment and optimisation of the chimera MerRCueR. **A)** Sequence alignment of MerR homologues. Both regulators MerR (*S. aureus*, “Sa”; accession code: CBI50741.1) and CueR (*E. coli*, “Ec”; accession code: CAD6020341) were aligned using the ClustalOmega tool^53^. The MerR derived DNA-Binding Domain (residues 1-38) is underlined blue, whilst the CueR derived C-terminal Domain (residues 37 – 135) is underlined orange. The fusion point (Arg37, CueR) is indicated in red with an arrow. Residues involved in Hg^2+^ coordination by MerR are indicated in dark blue, those involved in Cu^+^ coordination by CueR are indicated in orange. Note, Ser77 in CueR excludes divalent cations and is not involved in direct metal coordination with Cu^+^. Asterisk (*) indicates fully conserved residues, colon (:) indicates conserved residues with similar properties, and period (.) indicates residues of weakly similar properties. **B)** Homology model of the chimera MerRCueR. I-TASSER ^54^ was used to generated a homology of MerRCueR using the sequence from panel **(A)**. The top-ranking structural analogue was CueR from *E. coli* (PDB:1Q05; C-score = 0.71, TM-score= 0.904). The origin of each domain and their ligands are indicated using the same colour scheme. **C)** Circuit schematic comprising the designed chimera, MerRCueR. Bent arrows indicate promoters, flat-head arrows indicate inhibition, black semi-circles indicate ribosome binding sites, and genes for both *merRzntR* and *luxABCDE* coding sequences are indicated by the dark blue/orange and light blue arrows, respectively. **D)** Dose response of the chimera MerRCueR^WT^ following induction with a range of Cu^2+^ concentrations. **E)** Targeted mutagenesis of residues within loop region between α-helix 2 and α-helix. Amino acid substitutions were introduced into the parent hybrid gene *merRcueR* generating MerRCueR (MC) variants MC^A29T^, MC^A29T/G30P^ and MC^mut3^ (which carries the substitutions A29T/G30P/P32M). The dose response to CuSO_4_ is shown for all variants, with the limit of detection (LOD) shown only for MC^mut3^ (red) and EPA limit indicated. **F)** Substrate specificity of the MerRCueR circuit with variant MC^mut3^. Colours are indicated for the inducing metals only. For panels (**D-F**), cells were grown to OD_600_ =∼0.03 and induced with the concentrations of metal salts indicated, with luciferase activity (relative luminescence units [RLU]) normalised to optical density (OD_600_) values (RLU/OD_600_) measured and averaged for three time points (35, 40 and 45 mins) post-induction. Values are presented as mean and ± standard deviation of two or three independent replicates.

Exposure of this new strain to Cu^2+^ at a sublethal concentration of 1 mM failed to induce luciferase expression (Fig. 4D). As we had shown earlier that the structural interactions that couple the occupancy of the MBD to movement in the DBD were important for correct functioning of the MerRZntR chimera, we again inspected those residues in the homology model for MerRCueR. This revealed that several interactions between residues were absent in the MerRCueR hybrid ^17^ (Supplementary Fig. S4A-B). This included interactions between the side-chains of Glu46_MBD_ and Thr27_DBD_; the side chain of His43_MBD_ with the Pro28_DBD_ backbone ^17^; and the backbones of Thr38_MBD_ and Met30_DBD_. The corresponding residues at these positions in the *S. aureus* derived MerR_DBD_ are Ala29, Gly30 and Pro32, which would cause some disruption in the inter-domain communication network (Supplementary Fig. S4B). As residues Thr27_DBD_, Pro28_DBD_ and, to a lesser extent, Met30_DBD_ (a preference for a hydrophobic residue at this position) are highly conserved in CueR homologues from several genetic backgrounds (Supplementary Fig. S4C), we speculated that, as with the chimera MerRZntR, restoration of the hydrogen bonding network in MerRCueR to match that found natively in CueR would generate a functional copper responsive circuit.

To test this, we sequentially introduced amino acid substitutions A29T, A29T/G30P and A29T/G30P/P32M (strains SGB1028, SGB1029 and SGB1030, respectively). We observed gradual improvements in copper-responsive changes in activity from 1.1-fold with MerRCueR^WT^ to 64.3-fold with MerRCueR^A29T/G30P/P32M^ (from here on termed MerRCueR^mut3^ for simplicity) upon challenge with 1 mM Cu^2+^ (Fig. 4E). MerRCueR^mut3^ provided the highest RLU/OD_600_ output at each concentration and an LOD (10 μM) below the EPA guideline value of 20.45 μM (Fig. 4E). Moreover, the operational range was within the range of Cu^2+^ concentrations found in polluted environments ^57–59^, validating the potential use of this sensor as a relevant Cu^2+^ monitoring tool. These results confirmed that structure-guided mutagenesis indeed can be used to restore protein functionality following construction of MerR hybrid proteins. More generally, our results support the importance of residues relaying occupancy of the metal-binding site to movement in the DBD in metalloregulators^17,55^.

To assess the substrate-specificity, MerRCueR^mut3^ (SGB1030) was assayed in the presence of Cu^2+^, Hg^2+^, Cd^2+^, Pb^2+^, and Zn^2+^ as well as Ag^+^– a monovalent ion to which *E. coli* CueR is known to cross-react ^60,61^. Consistent with previous reports, MerRCueR^mut3^, did not show any response to divalent cations Hg^2+^, Cd^2+^, Pb^2+^, and Zn^2+^ (Fig. 4F). Cross-specificity was observed for Ag^+^, which revealed an LOD (0.01 μM) lower than values set by the EPA (0.46 μM) (Supplementary Fig. S5) and an operational range of the biosensor strain covering reported values for silver in polluted environments ^62^. This responsiveness to a monovalent metal ion is consistent with previous work ^61^, which showed that purified CueR protein in fact responds to monovalent Cu^+^ ions. In our experiments with living bacterial cells, copper was supplied as CuSO_4_ and thus Cu^2+^ ions. But uptake into the reducing conditions of the cytoplasm subsequently leads to conversion to Cu^+^, where this ion is detected by the CueR MBD, explaining the specificity profile of the biosensor of Cu^2+^ and Ag^+^.

Taken together, our results thus far highlighted that the modularity of the MerR regulators can be exploited to generate novel sensing modules, and a chimera-based strategy can be used to overcome species-specific design constraints such as promoter incompatibility. We envisage that a chimeric approach may be applicable to other protein families for import into a heterologous host such as *B. subtilis*, where using a closely related protein homologue from the desired host, or a related species, for example *S. aureus* could be a suitable donor of DNA-binding domains.

### MerR hybrids display preferences in promoter spacing distance

Past research has given detailed insights into how MerR regulators bind and differentiate between metal ions ^63–67^, which allowed us to develop the functional hybrid regulators described above. However, to fully engineer these proteins for synthetic biology applications, detailed knowledge is also required of the factors that enable MerR regulators to correctly under-twist their respective cognate promoters. MerR target promoters generally possess either a 19- or 20-bp spacer between the -10 and -35 elements. In the following text, a MerR homologue will be denoted with a subscript of spacing in its target promoter, for example CueR_19_ acts upon the P_*copA19*_ promoter, which possesses a spacer region of 19 bp. Perturbation of this spacer region in MerR family promoters is known to disrupt correct transcriptional regulation of the promoter ^68^. Interestingly, previous work on ZntR_20_ has suggested that the MBD, rather than the DBD, determines the degree of promoter distortion ^23^. This would imply that regulators CueR_19_ and ZntR_20_, which act on promoters with 19- and 20-bp spacer regions, respectively, would distort their respective promoters to different degrees, based on their MBDs, as previously suggested by Brown *et al*. ^15^. This may have implications for chimeric MerR transcription factors that could prefer a target promoter whose spacer region length is determined by the origin of the MBD, even if the same DBD was used in all chimeras. For example, in the MerRCueR^mut3^ regulator we described above, we had so far used the 20 bp-spaced promoter P_*merR20*_ to drive expression of the luciferase reporter. This promoter is natively recognised by MerR_20_ of *S. aureus*. However, the donor protein of the MBD, *E. coli* CueR_19_, natively controls a 19 bp-spaced promoter, P_*copA19*_. If it is indeed the MBD that determines promoter distortion, we might hypothesise that the MerRCueR^mut3^ chimera should perform better when provided with a promoter with a 19-bp spacer region.

To test the effects of the C-terminal MBD region of our hybrid proteins on promoter regulation, we investigated the ability of MerR and MerRCueR^mut3^ to regulate the activity of either a 19- or 20-bp spaced promoter. P_*merR20*_ is the *S. aureus* promoter that we have used so far, and P_*merR19*_ is its derivative in which 1bp was deleted (Supplementary. Fig. S6). When MerR_20_ was used to regulate the activity of the P_*merR19*_*-lux reporter*, the dynamic range of induction was severely perturbed, with higher basal activity when compared to P_*merR20*_. (Fig. 5). This was consistent with previous studies of the *mer* operon of *Tn501* in Gram-negative systems, where increased basal expression was seen when the 19-bp spacer was shortened ^68^. Thus, MerR of *S. aureus* worked better when provided with its native promoter in the heterologous *B. subtilis* system, as expected.

**Figure 5.**
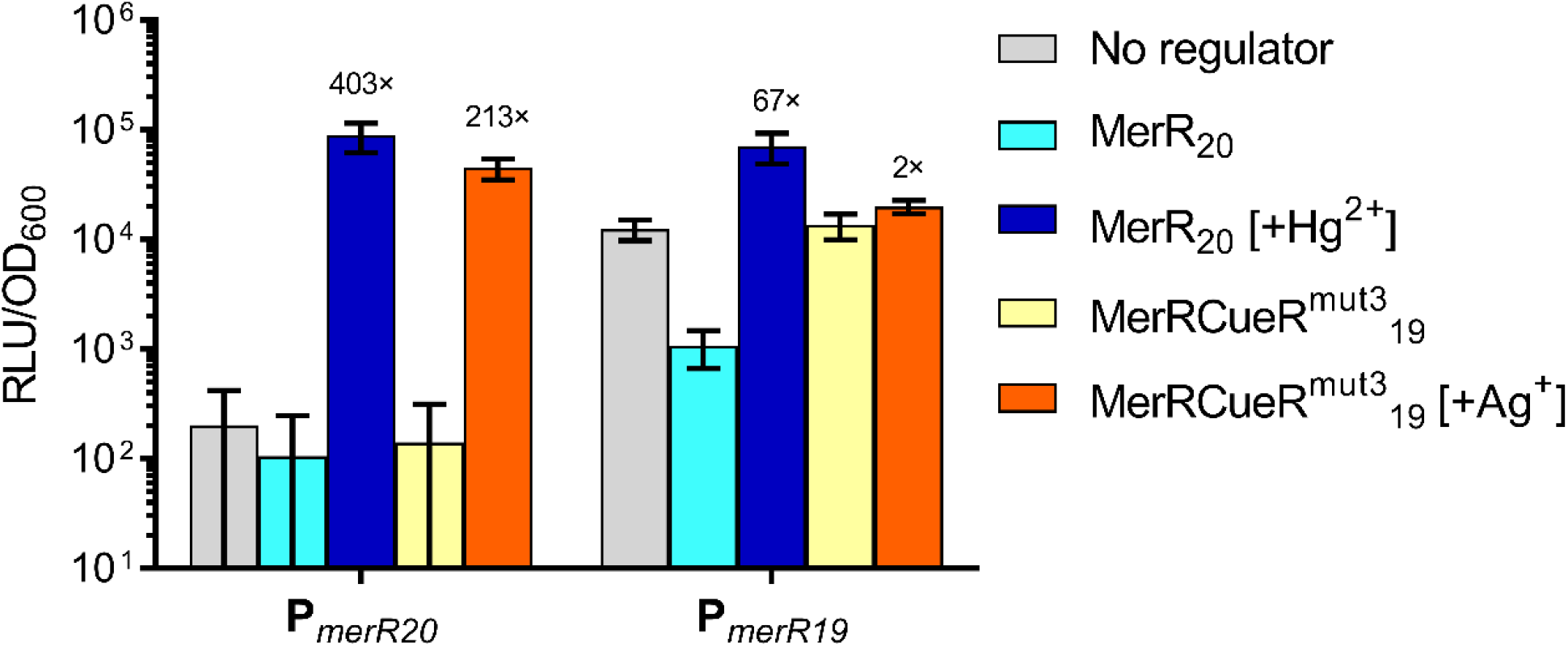
Regulation of promoters P_*merR20*_ and P_*merR19*_ by wild-type MerR and the chimeric MerRCueR^mut3^. The activity of either promoters P_*merR20*_ and P_*merR19*_ fused to the luciferase reporter was tested in either the absence of any regulator (grey), or in the presence of the regulator MerR (blue) or MerRCueR^mut3^_19_ (yellow/orange). The experiments were carried out in the absence (lighter colours) or presence of an inducing metal (MerR, dark blue [100 nM Hg^2+^], MerRCueR^mut3^ _19_, orange [3 μM Ag^+^]). Cells were grown to OD_600_ =∼0.03 and then induced with the metals as indicated, with luciferase activity output (relative luminescence units [RLU]) normalised to optical density (OD_600_) values (RLU/OD_600_) measured and averaged for three time points (35, 40 and 45 mins) post-induction. Fold values represent the induction ratio between induced against uninduced. Values are presented as mean ± standard deviation of two or three independent replicates.

Next, we compared the ability of MerRCueR^mut3^ to regulate the activity of both P_*merR20*_ and P_*merR19*_ transcriptional reporters. Surprisingly, in the system containing the P_*merR19*_ promoter, the presence of the MerRCueR^mut3^ regulator did not lead to any change in promoter activity compared to cells lacking a regulator. This was not changed whether the inducer (Ag^+^) was present or not, suggesting the chimeric protein was unable to interact with the shorter spaced target promoter. In contrast, the chimeric regulator was able to elicit a 213-fold increase in the promoter activity of P_*merR20*_ when challenged with Ag^+^ (Fig. 5). This strongly suggests that in the MerRCueR^mut3^ hybrid, the MBD does not determine optimal promoter spacing, contrary to earlier reports on ZntR ^23^. It is currently not clear whether this is because the DBD and promoter were derived from a Gram-positive system, where the DBD may play a more important role in determining promoter spacing, or whether the earlier reports on ZntR were a particular feature of that specific system. Different combinations of MBD and DBDs would need to be tested using different output modules to answer this. However, we can conclude that for the *B. subtilis* system used here, we appear to be able to construct and use chimeric regulators with diverse metal specificity without the need of adjusting promoter spacing in the output module.

### The natively split sigma factor SigO-RsoA enables the design of modular AND logic circuits

Having demonstrated the utility of and some design rules for chimeric regulators as novel metal-responsive circuits, we wanted to test whether we could overcome possible problems with cross-specificity between metals by designing a modular AND logic gate. This type of genetic gate requires the presence of two inputs in order to produce an output ^41^. To illustrate, none of our metal biosensors responded to only a single metal. But it may be possible to use AND logic to combine two different metal sensors whose substrate specificity overlaps only for one metal. In such a case, this metal would be the only substrate to trigger both regulators, and therefore the output signal would be produced in response only to this single metal, creating a highly specific biosensor. A similar approach has already proved effective in generating an ultra-specific metal sensor circuit in *E. coli* ^9^. However, a standardised, easy to use two-input AND gate system, offering modular assembly and fast response times, is currently not available in the suite of genetic toolboxes for *B. subtilis*.

Therefore, to generate such a system, we exploited the *B. subtilis* natively split-sigma factor system SigO-RsoA. Various studies have demonstrated that both SigO and RsoA, which constitute domains σ^4^ and σ^2^ of the sigma factor, respectively, must co-operate to initiate transcription from the promoter P_*oxdC*_ ^69,70^. This system thus effectively acts as a natural biological AND gate, which should be amenable for engineering such regulatory logic in synthetic circuits. To facilitate fast assembly of the AND gate, we generated a series of new plasmids termed SANDBOX (*<u>S</u>ubtilis* <u>AND BOX</u>) based on the Golden Gate assembly format using the type II restriction enzyme BsaI. This toolbox includes vectors pBSAND1 (carrying *rsoA*), pBSAND2 (carrying *sigO*), pBSANDlux (carrying the P_*oxdC -*_*luxABCDE* reporter), and a CRISPR based deletion plasmid (pBSANDdel) used to delete the native SigO-RsoA divergent regulon^71^. The architecture of all the vectors and the Golden Gate cloning site sequences can be found in Supplementary Fig. S7.

For the design and validation of our initial two-input AND gate, we used two well-characterised *B. subtilis* promoters, P_*xylA*_ and P_*liaI*_ ^31^, to control transcription of SigO and RsoA, respectively. The output from the gate (luciferase activity) was driven by the cognate promoter of this split σ-factor system, P_*oxdC*_. The overall architecture of the circuit is shown in Fig. 6A. To assess the functionality and determine optimum induction of the AND gate, we assayed different combinations of bacitracin (P_*liaI*_-inducer) and xylose (P_*xylA*_-inducer) concentrations, revealing the system behaved correctly in an AND gate manner ^41^, i.e. reporter induction was only observed if both inducers were added together. We did observe some leakiness from P_*xylA*_, giving weak outputs when bacitracin alone was used, which was consistent with a previous study that characterised the activity of this promoter ^31^. Overall, our data demonstrated that the new standardised logic system was functional in *B. subtilis* and should allow for flexible and rapid assembly of two-input AND gate circuits for a variety of applications.

**Figure 6.**
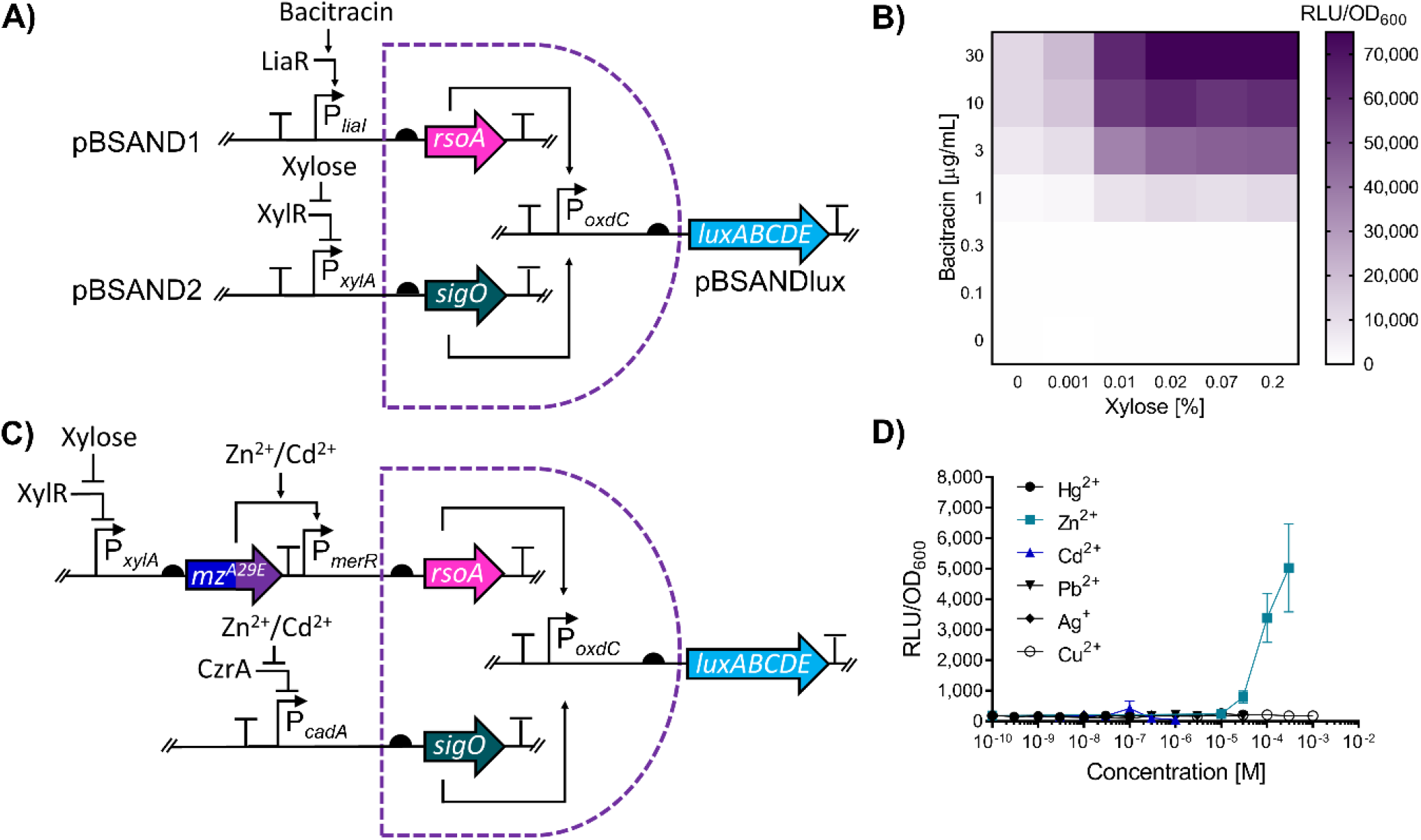
Assessing the functionality and modularity of the split SigO-RsoA σ-factor two-input AND gate for *B. subtilis*. **A)** Circuit schematic of the two-input AND gate. To test the functionality of the SigO-RsoA based AND gate, two promoters P_*liaI*_ and P_*xylA*_, inducible by bacitracin and xylose, respectively, were used to induce the expression of both SigO and RsoA, allowing for transcription from the P_*oxdC*_ promoter to drive luciferase (*luxABCDE*) expression. The *sigO* and *rsoA* genes are indicated via the teal and pink arrows respectively. **B)** Functionality of the SigO-RsoA AND gate using bacitracin and xylose. Cells with the integrated SANDBOX vectors incorporating a bacitracin and xylose inducible promoter were induced with various combinations of xylose and bacitracin with the heatmap used to show luciferase output across the tested concentrations. **C)** Circuit schematic for an ultra-specific Zn^2+^ biosensor used the SigO-RsoA system. SigO and RsoA genes are indicated as previously described. The circuit, which utilises the chimera MerRZntR^A29E^ is denoted via the split dark blue and purple arrow and simplified to “*mz*^*A29E*^*”*.. For simplicity, the native *B. subtilis* metalloregulator CzrA present at a different genomic locus on the chromosome has not been indicated. **D)** Dose response and specificity of the SigO-RsoA based Zn^2+^ detection circuit. For panels **(B)** and **(D)**, cells were grown to OD_600_ =∼0.03 and induced with the concentrations of either xylose and bacitracin **(B)** or metals **(D)** as indicated. Luciferase activity output (relative luminescence units [RLU]) was normalised to optical density (OD_600_) values (RLU/OD_600_) measured and averaged for three time points (35, 40 and 45 mins) post-induction. Values are presented as mean ± standard deviation of two or three independent replicates.

Having confirmed the functionality of our SANDBOX system, we sought to combine it with our chimeric biological parts for metal detection to develop an ultra-specific zinc biosensor. As mentioned above, this required two different metal-inducible regulators with overlapping substrate specificity for one metal. Moreover, the AND gate requires the use of two target promoters without cross-recognition by the two regulators. Given our MerR-based systems all converge on the same output promoter, we had to source the second component of our AND gate system from an unrelated metal sensor. Therefore, we decided to use the optimised chimera MerRZntR^A29E^, together with CzrA – a native metalloregulator in *B. subtilis* belonging to the ArsR protein family ^72^. Whilst we had already determined the specificity profile for MerRZntR^A29E^ (Fig. 3C), we needed to determine this for CzrA. For this, we generated a transcriptional luciferase reporter of the CzrA target promoter P_*cadA*_ and exposed cells harbouring this reporter to the same panel of metals used before. This revealed induction of P_*cadA*_-*luxABCDE* in the presence of Zn^2+^ and, to a minor degree, Cd^2+^ (Supplementary Fig. S8). Thus, both MerRZntR^A29E^ and CzrA responded strongly to Zn^2+^ but shared no other substrates. They should therefore be a suitable pair for construction of the AND gate biosensor. To note, whilst both regulators responded to Cd^2+^, transcriptional output from P_*cadA*_ was negligible compared to the output produced from MerRZntR^A29E^ (Fig. 4C and Supplementary Fig. S8 and thus unlikely to create interference.

Based on this information, we proceeded to construct the AND gate-based Zn^2+^-specific circuit, shown schematically in Fig. 6C. Expression of SigO was placed under control of P_*cadA*_ (CzrA), and RsoA expression was placed under control of P_*merR*_ (MerRZntR^A29E^). To generate a detectable output, luciferase expression was again controlled from P_*oxdC*_ (SigO-RsoA). When the strain harbouring this genetic circuit was tested against the same panel of metals as above, a strong response was only obtained in the presence of Zn^2+^, showing a clear dose-response behaviour (Fig. 6D). As anticipated, the signal elicited by the presence of Cd^2+^ was very low and barely detectable above background. This showed that the use of a simple logic AND gate significantly reduced the noise generated from non-target metals, such as Cd^2+^, to generate a highly specific Zn^2+^ biosensor from individual regulatory parts that each respond to multiple different metals. Furthermore, the resulting data confirmed that the *B. subtilis* SigO-RsoA system can be exploited for the design of robust AND gates, with the modularity of the system demonstrated through the adaptation of this system in the design of an ultra-specific biosensor circuit.

## CONCLUSIONS

Monitoring of environmental levels of heavy metal contamination is integral in the management of the risk associated with polluted areas and to assess the efficacy of remediation efforts at these sites. Whole cell biosensors offer an attractive alternative to conventional monitoring methods, which can be expensive and resource heavy. In this work, we demonstrated that novel hybrid transcription factors can be assembled using the modular MerR proteins from different bacteria, and that their functionality as biosensors in *B. subtilis* can be optimised using structure-guided mutagenesis. This approach will allow researchers to tap into the great diversity of substrate specificity found in the MerR proteins from Gram-negative bacteria for use in Gram-positive chassis organisms, by utilising their metal-binding domains in a hybrid protein that has a constant DNA-binding domain from a Gram-positive donor. This is important in overcoming problems commonly faced when sourcing genetic components from different species for use in established chassis systems such as *B. subtilis*, including issues with promoter recognition and compatibility with the host’s transcription machinery.

We initially demonstrated the functionality of a heterologous MerR based regulatory circuit from *S. aureus* for the detection of Hg^2+^ ions in *B. subtilis* and showed that the modular architecture of MerR can be exploited to generate novel chimeric regulators by replacing the metal-binding domain of one regulator with one from another protein with different specificity. Maintaining the same DNA-binding domain in all proteins addresses problems of promoter recognition. We further demonstrated that the functionality and sensitivity of these circuits can be improved through structure-guided design, allowing monitoring of metal contamination in environmentally relevant ranges. While some chimeras with a MBD from Gram-negative-derived proteins may not immediately be functional, we have shown here how engineering of residues in the communication interface between MBD and DBD can be used to restore function.

Finally, we showed that problems with cross-specificity can be resolved by incorporating our novel orthologous regulators into AND gate-based logic circuits that include native *B. subtilis* metalloregulators. For this, we utilised the *B. subtilis* natively split σ-factor system SigO-RsoA and demonstrated that this can drastically reduce signal from non-target contaminants. Initial construction and validation of the circuit was done using well-characterised *B. subtilis* promoters, with subsequent demonstration of the modularity of this system in the design of an ultra-specific Zn^2+^ sensor based on overlapping specificities of two metalloregulators.

Apart from the design and testing of novel biosensors, this work has led to the development of a new toolbox of Golden Gate-based vectors, which enables easy construction of two-input AND gates in *B. subtilis*. We anticipate that based on the modularity of this system, it will not only be useful for the design of a variety of biomonitoring tools, but also can be adopted for a range of applications such as biomedical diagnostics or metabolic engineering. Taken together, this work provides insights into how modular regulators, such as the MerR family, can be exploited in the design of synthetic circuits for the detection of heavy metal contaminants. We have shown how structure-guided design can produce functional sensors even when protein domains are sourced from distinct species, which may also inform work on other proteins with similar modular architecture.

## MATERIALS AND METHODS

### Bacterial Stains, Growth Conditions and reagents

All strains used in this study are listed in Table S1 and were routinely grown in Lysogeny Broth (LB; 10 g/L tryptone, 5 g/L yeast extract, 5 g/L NaCl) at 37°C with aeration (agitation at 180 rpm). Solid media contained 1.5 % (wt/vol) agar. Selective media for *B. subtilis* contained chloramphenicol (5 μg/ml) or spectinomycin (100 μg/ml), whilst selective media for *E. coli* contained ampicillin (100 μg/ml). For luciferase assays, *B. subtilis* strains were grown in a modified M9 minimal media (MM9). The composition of MM9 was as follows: 1 mM MgSO_4_, 0.3% fructose, 1% casamino acids, 0.05 mM FeCl_3_/ 0.1 mM citric acid solution, deionised water and 1x M9 salts (31.7 mM Na_2_HPO_4_, 17.22 mM K_2_HPO_4_, 17.11 mM NaCl, 9.34 mM NH_4_Cl). For transformations of *E. coli* DH5α (see below), SOC medium was used with the following composition: 2% tryptone, 0.5% yeast extract, 10 mM NaCl, 2.5 mM KCl, 10 mM MgCl_2_, 10 mM MgSO_4_, and 20 mM glucose. For transformations of *B. subtilis*, MNGE medium was used based on the composition described by Radeck *et al* ^31^. Further details about the transformation procedure can be found below. Metal salts Ag(NO_3_), ZnSO_4_ .7H_2_O, Pb(NO_3_) and CdCl_2_ were obtained from Fisher Scientific, CuSO_4_7H_2_O and HgCl_2_ were obtained from Sigma-Aldrich.

### DNA manipulation, plasmid construction and bacterial transformation

Detailed information regarding strains, plasmids, primers, and genetic sequences for biological parts used in this used in this study are listed in supplementary materials, as tables S1, S2, S3 and S4, respectively. All cloning steps, including restriction endonuclease digestion, ligation and PCRs, used enzymes and buffers from New England Biolabs (NEB; Ipswich, MA, USA) according to the relevant NEB protocols. All PCR clean-up kits (Monarch® PCR clean-up kit), plasmid (Monarch® PCR mini-prep kit) and gel extraction kits (Monarch® DNA gel extraction kit) were also obtained from NEB and used according to the manufacturer’s protocols. For ligation of inserts into plasmid vectors used, except for the Golden Gate procedure described below, the NEB T4 DNA ligase protocol M0202 was used. All PCR amplifications were performed using Q5® DNA polymerase (NEB protocol M0491), whereas for colony PCR to confirm ligation of inserts into desired vectors, OneTaq® polymerase was used (NEB M0480). Chemically competent DH5α cells were transformed with isolated plasmids or ligation reactions using a heat-shock procedure in which cells were mixed with DNA for 10 minutes on ice, heat-shocked at 42 °C for 90 seconds, placed back on ice for 5 minutes after which SOC medium was added, and cells were incubated at 37 °C in a shaking incubator (200 rpm) for 1 hour before plating onto selective media (see above). Transformations of *B. subtilis* were carried out as described by Harwood and Cutting^73^ with integration of plasmids derived from pAH328 at the *sacA* locus confirmed using colony PCR with *sacA* up- and down-primers SG0528/SG0529 and SG0530/SG0531, respectively, and integration of pXT-derived plasmids at the *thrC* locus confirmed via threonine auxotrophy, as described by Radeck *et al* ^31^.

### Plasmid construction

To amplify P_*merR*_ *and* MerR, *S. aureus* TW20 genomic DNA (gDNA) was isolated using a GeneJet genomic DNA extraction kit (Thermofisher). P_*merR20*_ and MerR were amplified using primers SG0985/SG0986 and SG0987/SG0988, respectively, with 20 ng of *S. aureus* gDNA as a template. The amplified promoter and regulator were digested with EcoRI-/SpeI-HF and BamHI-/EcoRI-HF, respectively, and ligated into vectors pAH328 and pXT, respectively to generate plasmids pJGlux01 and pJGXT01, respectively. To generate a 1-bp deletion of the P_*merR20*_ promoter (P_*merR19*_), we used site-directed mutagenesis with primers designed as described by Liu and Naismith^74^, and amplification performed using the Q5® DNA polymerase protocol as described above using 20 ng of pJGlux01 as a template and 2.5 nM of primers SG1124/SG1125 (50 μL reaction). For this, 12 amplifications cycles, an extension time of 1 minute per kb, and an annealing temperature of 60 – 66 °C were used. Following amplification, DpnI was added directly to the reaction to a final concentration of 400 U/mL, with amplification confirmed using agarose gel electrophoresis. This generated plasmid pJGlux02. To generate the P_*veg*_ -luciferase fusion, the promoter was excised from pSB1C3-P*veg* ^31^ using EcoRI-/SpeI-HF and ligated into EcoRI-/SpeI-HF digested pAH328 to generate plasmid pJGlux03. To generate the promoter fusion for Gram-negative-derived MerR (P_*cadA19*_), primers SG1164/SG1171 were used to amplify the P_*cadA19*_ promoter from *Pseudomonas aeruginosa* PAO1 gDNA (20 ng). The amplified promoter was digested with EcoRI/SpeI-HF and ligated into pAH328 to generate pJGlux04.

### Metal sensitive chimeras

To construct chimeric regulator MerRZntR, the MerRZntR amino acid sequence was designed on a previous chimeric regulator as described by Brocklehurst *et al* ^23^ using residues Met1-Tyr38 of the MerR_DBD_ region and residues Arg38 (used as a junction point between both proteins) to Cys141 of ZntR of *E. coli* MG1655. The resulting chimeric DNA sequence was flanked a 3’-BamHI and 5’-EcoRI sites and commercially synthesised (GenScript, Rijswijk, Netherlands) into vector pUC19 (pJGUC01). The insert was excised with BamHI-/EcoRI-HF (1 μg) and sub-cloned into BamHI-/EcoRI-HF digested pXT to generate pJGXT02. To generate variants MerRZntR^A29E^, MerRZntR^A29E/G30H^ and MerRZntR^A29E/G30H/P32V^, site-directed mutagenesis approach was used as described above using mutagenic primer pairs SG1172/SG1173, SG1174/SG1175 and SG1200/SG1201, respectively, which generated plasmids pXTJG15, pXTJG16 and pXTJG23, respectively.

The chimeric regulator MerRCueR was constructed in a manner analogous to that of MerRZntR using amino acid residues Met1 – Tyr38 of the MerR DBD (*S. aureus*) and residues Arg37 (used as a junction point) to Gly135 of CueR of *E. coli* MG1655. The resulting chimeric DNA sequence was flanked with the previously mentioned restriction sites as for MerRZntR and synthesised into pUC19 (pJGUC02). MerRCueR was subsequently subcloned into pXT as described above for the generation of pJGXT02 (see above), with the resulting plasmid designated pJGXT03. For the construction of MerRCueR^A29T^ and MerRCueR^A29T/G30P^, site-directed mutagenesis was performed as described above using plasmid pJGXT03 as template with mutagenic primers SG1142/SG1143 and SG1154/SG1155 to generate plasmids pJGXT07 and pJGXT08, respectively. For MerRCueR^A29T/G30P/P32M^ (MerRCueR^mut3^), plasmid pJGXT08 was used as a template with primers SG1167/SG1168 to generate plasmid pXTJG11.

### The Bacillus SANDBOX Plasmids p

#### BSAND1

The *Bacillus* BioBrick vector pBS4S ^31^ was used as a parent plasmid for the construction of pBSAND1. The BsaI site in the *bla (*amp^r^) gene of pBS4S was removed using primer pairs SG1242/SG1243 to generate plasmid pJGBS4S01. The *rfp* cassette was amplified from pBS4S using primers SG1272/SG1273 to incorporate a 5’-BsaI site and 5’-terminator sequence and 3’-BsaI and 3’-SfiI site. *rsoA* was amplified from *B. subtilis* W168 gDNA (20 ng) using primers SG1275/ SG1250 to incorporate a 5’-SfiI site and a 3’-terminator and 3’-PstI site. Amplified *rfp* and *rsoA* were digested with EcoRI-/SfiI-HF and SfiI-/PstI-HF, respectively, and ligated into EcoRI-/PstI-HF digested pJGBS4S01 in a single reaction. The removal of BsaI from *bla*, and the insertion of the new BsaI sites flanking RFP were confirmed by restriction digestion with BsaI, and the ligation of both *rfp* and *rsoA* into pJGBS4S01 confirmed using PCR with primers SG601/SG602 and Sanger sequencing (Eurofins Genomics, Germany). The resulting plasmid allowing for expression of RsoA from a promoter of choice was designated pBSAND1 and can be linearised for transformation of *B. subtilis* using the enzyme ScaI. The terminator sequence used is an *in silico* designed terminator called “Term 1” ^75^, whilst to ensure strong expression of RsoA, the RBS sequence R1 described by Guiziou *et al* ^76^ was used. The resulting plasmid was designated pBSAND1 and can be linearised using ScaI to allow for integration in *B. subtilis* at *thrC*.

For plasmid pBSAND1-P_*liaI*_, *P*_*liaI*_ was amplified from *B. subtilis* gDNA using primers SG1388/SG1389 and assembled into pBSAND1 using Golden Gate cloning (see below) to allow for bacitracin inducible expression of RsoA. To construct pBSAND1-*P*_*xylA*_-MerRZntR^A29E^-Terminator-P_*merR20*_, *P*_*xylA*_ was amplified from pSB1A3-P_*xylA*_ ^31^ using primers SG1297/SG1410; MerRZntR^A29E^ (including its native RBS) was amplified from pJGXT15 using primers SG1382/SG1383; and *P*_*merR20*_ (to include a 5’ terminator, “Term 1”) was amplified from pJGlux01 using primers SG1384/1385. The three fragments were assembled into pBSAND1 using Golden Gate to allow for metal-inducible expression of RsoA.

#### pBSAND2

The *Bacillus* BioBrick vector pBS2E^31^ was used as a parent plasmid for the construction of pBSAND2. The BsaI site in the *bla* gene of pBS2E was removed using SG1242/SG1243 generating pJGBS2E03, after which an NgoMIV site was inserted in the *bla* gene using primers SG1245/SG1246, generating plasmid pJGBS2E04. The same *rfp* cassette was used as for pBSAND1, whilst *sigO* was amplified from *B. subtilis* gDNA using primers SG1274/SG1248 to incorporate a 5’-SfiI site, a 3’-terminator and a 3’-PstI site. Both amplified *rfp* and *sigO* were digested with EcoRI-/SfiI-HF and SfiI-/PstI-HF, respectively and ligated into EcoRI-/PstI-HF digested pJGBS2E04. Removal of BsaI from *bla*, the insertion of NgoMIV into *bla*, and the insertion of new BsaI sites flanking RFP were confirmed by restriction digestion with BsaI and NgoMIV. Ligation of *rfp* and *sigO* into pJGBS2E04 was confirmed by PCR using primers SG0245/SG0246 and sequencing (Eurofins Genomics, Germany). The resulting plasmid was designated pBSAND2 and can be linearised using NgoMIV to allow for integration in *B. subtilis* at *lacA*.

For plasmid pBSAND2-P_*xylA*_, P_*xylA*_ was amplified from pSB1A3-P_*xylA*_ ^31^ using primers SG1297/SG1298 and assembled into pBSAND2 using Golden Gate to allow for xylose inducible expression of SigO. For pBSAND2-*P*_*cadA*_, P_*cadA*_ was amplified from *B. subtilis* gDNA using primers SG1403/SG1387 and assembled into pBSAND2 using Golden Gate (see below) to allow for metal-inducible expression of SigO

#### pBSANDlux

The *Bacillus* BioBrick vector pBS3Slux was used as a parent plasmid for the construction of pBSANDlux (P_*oxdC*_*-luxABCDE* reporter). The BsaI sites in the *luxC* and *bla* genes were removed via site-directed mutagenesis as described above using primer pairs SG1239/SG1240 and SG1242/SG1243, respectively, to generate plasmids pJGBS3Clux02 and pJGBS3Clux03, respectively. The *rfp* cassette was amplified from pBS4S using primers SG1272/SG1324, to incorporate a 5’ terminator as well as flanking 5’- and 3’-BsaI sites which was subsequently digested using EcoRI/PstI and ligated into EcoRI-/PstI-HF digested pJGBS3Clux03. To confirm the presence of the *rfp* cassette with a 5’-terminator, colony PCR was performed primers SG1303/SG1325 and constructs sequenced using SG0991. Removal of BsaI sites in *bla* and *luxC*, as well as the incorporation of BsaI sites flanking RFP were confirmed using restriction digestion using BsaI. The resulting plasmid pBSGGlux can be linearised using ScaI for integration in *B. subtilis* at the *sacA* locus.

Finally for plasmid pBSANDlux, required as part of the *Bacillus* SANDBOX system to generate 2-input biosensors, P_*oxdC*_ was amplified using primers SG1299/SG1300 from *B. subtilis* W168 gDNA (20 ng) and assembled into pBSGGlux using Golden Gate (see below). When utilised with plasmids pBSAND1-P_*liaI*_ and pBSAND2-P_*xylA*_, pBSANDlux allows for bacitracin and xylose inducible luciferase output. When utilised with pBSAND1-P_*xylA*_-MerRZntR^A29E^-Terminator-P_*merR*_, and pBSAND2-P_*cadA*_, pBSANDlux allows for metal-inducible luciferase output.

#### pBSANDdel

The CRISPR-Cas9 deletion plasmid pJOE8999 was used as the parent plasmid for the construction of pBSANDdel, with construction done as described by Altenbuchner ^77^. To amplify left and right homology regions surrounding the *sigO-rsoA* operon, primers SG1326/1327 and SG1328/SG1329 were used. These fragments were digested with SfiI-HF and ligated into SfiI-HF digested pJOE8999. The insertion of the flanking homology regions was confirmed using colony PCR with primers SG1347/SG1348. To insert the gRNA to direct the Cas9 machinery, 5 μL of oligonucleotides SG1349/SG1350 (100 μM) were mixed with 90 μL of 30 mM HEPES (pH 7.8), heated to 95 °C for 5 mins and then cooled to 4 °C at a rate of 0.1 °C/sec. The annealed oligos (2 μL) were ligated into pJOE8999 containing the left/right homology regions using Golden Gate and the incorporation of the gRNA confirmed using blue-white screening on LB agar supplemented with X-Gal (50 μg/mL). The resulting vector pBSANDdel cuts upstream of the gene *rsoI* and allows for removal of the entire *sigO-rsoA* operon in *B. subtilis* W168.

### Golden Gate Assembly

For the cloning of inserts using Golden Gate, the reaction (10 μL) comprised final concentrations of 10,000 U/mL T4 DNA ligase, 1,000 U/mL BsaI-HFv2, 1× T4 DNA ligase buffer, 1× CutSmart Buffer, 100 ng of plasmid and equimolar amounts of inserts. Note for the assembly of pBSAND1-P_*xylA*_-MerRZntR^A29E^-Terminator-P_*merR*_, a 3:1 insert to vector ratio was used. The reaction conditions were as follows, 37°C for 5 mins, 5-10 cycles of 37°C for 5 mins and 16°C for 10 mins, followed by 16 °C for 30 mins, 50 °C for 5 mins and 80 °C for 10 mins. For assembling three or more inserts into a vector, we found increasing the numbers of cycles to 30 or more beneficial.

### Homology modelling & structural analysis

Homology models for both MerRZntR and MerRCueR were generated using the online I-TASSER server with default parameters, with the respective homology model with the highest C-score selected for further analysis ^54^. As only partially resolved structures for ZntR are available in the Protein Data Bank (all of which lack residues 1-68, PDB: 1Q09), a homology model of a full-length ZntR monomer was also constructed using the amino acid sequence of ZntR from *E coli* MG1655. For all generated homology models, the closest structural analogue was selected and used as a structural reference for quality control. For models of ZntR, MerRZntR and MerRCueR, the C-scores were 0.71 for all the generated models, all of which shared the same closest structural analogue (CueR, PDB:1Q05) as determined by I-TASSER. The TM-structural alignment programme within I-TASSER compared closest structural analogue, PDB 1Q05, to all the generated structures of ZntR, MerRZntR and MerRCueR with TM-scores of 0.71, 0.71, 0.81 respectively, where a score of > 0.5 indicates a similar fold. For visualisation of all structures, the resulting PDB structures generated by I-TASSER were imported into PyMol Version 2.0 (Schrödinger, LLC) for visualisation ^78^.

### Luciferase assays

For luciferase reporter assays, overnight cultures of each strain to be tested were inoculated 1:1000 into the modified M9 minimal medium described above with added xylose (0.2% final concentration) and distributed into 96-well microplates (Corning; black, clear, flat bottom), with 95 μL culture volume per well. Wells around the plate edge were filled with water to minimise evaporation. A Tecan Spark microplate reader (Tecan Trading AG, Switzerland) was used to monitor luciferase activity and OD_600_ values for each strain. Cells were grown with continuous shaking incubation (37°C; 180 rpm; orbital motion; amplitude, 3 mm) until OD_600_ = ∼0.03 (corresponding to OD_600_ = ∼0.3 when measured in cuvettes of 1 cm light path length), after which cells were induced with 5 μL of metal stock solutions to reach varying final concentrations (100 μl final volume). For metal induction experiments, the following metal salts were used: HgCl_2_, ZnSO_4_, CuSO_4_, CdCl_2_, Pb(NO)_3_, and Ag(NO)_3_. Measurements of OD_600_ and luminescence (relative luminescence units [RLU]) were measured every 5 minutes for 120 minutes. RLU and OD_600_ values were blank-corrected using the average of triplicate measurements of RLU and OD_600_ for MM9 medium alone. Luminescence activity was normalised to cell density for each time data point and reported as RLU/OD_600_. For all dose-response and metal-specificity studies, final RLU/OD_600_ values were the average of three time points (35, 40, and 45 min) after challenge with heavy metal salts. All experiments were performed in biological triplicates. All data were processed using Microsoft Excel and subsequently analysed in GraphPad Prism 7. To determine the Limit of Detection (LOD) for our sensors we followed the methods as described by Armbruster and Pry ^79^ and Wan *et al* ^8^.

## Data accessibility

All data used to generate the figures will be made available in additional file 2

## Supporting information

Additional file 1: Supplementary Tables and Figures (DOCX)

Additional file 2: Figure data. (XLSX)

## ACKNOWLEDGEMENTS

We acknowledge the Engineering and Physical Sciences Research Council (EPSRC; EP/PO2081X/1) and industrial collaborators/partners for funding the Resilient Materials for Life (RM4L) project. J.G. was supported by a University of Bath Leveraged Studentship Award. We thank the technical staff in the Life Sciences Department for key support.

## AUTHOR INFORMATION

### Corresponding Authors

**Bianca Jean Reeksting**, Department of Life Science, University of Bath, Bath BA2 7AY, United Kingdom.

**Susanne Gebhard**, Department of Life Science, University of Bath, Bath BA2 7AY, United Kingdom.

### Authors

**Jasdeep Singh Ghataora**, Department of Life Science, University of Bath, Bath BA2 7AY, United Kingdom

### Author Contributions

BJR, JSG, and SG conceived the study; JG conducted all experimental work and data analysis; SG acquired funding, BJR and SG coordinated the work; JG, BJR and SG wrote the manuscript.

