## Supplementary Tables and Figures for "Chimeric MerR-Family Regulators and Logic Elements for the Design of Metal Sensitive Genetic Circuits in *Bacillus subtilis*"


**Supplementary Table S1**. Strains used within this study 2

**Supplementary Table S2**. Plasmids used in this study 4

**Supplementary Table S3**. Primers used in this study 7

**Supplementary Table S4**. Genetic Sequences used in this study 11

**Supplementary Figure S1**. Comparison of Gram +ve and Gram -ve MerR promoter activity in *B. subtilis* 15

**Supplementary Figure S2**. Comparison of ZntR sequences and structural analysis of the chimera MerRZntR 16

**Supplementary Figure S3**. Activity of double and triple MerRZntR mutants against the wild-type and single mutant MerRZntRA29E 17

**Supplementary Figure S4**. Comparison of inter-domain communication between CueR and the chimera MerRCueR 18

**Supplementary Figure S5**. Dose response of $P_{merR}$ regulated by MerRCueRmut3 in response to $Ag^{+}$ induction 19

**Supplementary Figure S6**. Wild-type ($P_{merR20}$) and mutant ($P_{merR19}$) promoters 20

**Supplementary Figure S7**. *The B. subtilis* SANDBOX plasmids 21

**Supplementary Figure S8**. *Bacillus subtilis* natively regulated CzrA based circuit 22

**References** 23

Supplementary Table S1. Strains used within this study

| Strain | Characteristics[a] | Construction[b] |
|---|---|---|
| DH5α | *Escherichia coli* DH5α | Laboratory stock |
| W168 | Wild-type *Bacillus subtilis* W168 | Laboratory stock |
| PAO1 | Wild-type *Pseudomonas aeruginosa* PAO1 | Laboratory stock |
| SGB1003 | W168 *sacA*:: pJGlux01-P$_{merR20}$-*luxABCDE-cm*$^{r}$ | pJGlux01 -> W168 |
| SGB1005 | W168 *sacA*:: pJGlux01-P$_{merR20}$-*luxABCDE-cm*$^{r}$; *thrC*::pJGXT01- *P*$_{xylA}$-*MerR- spc*$^{r}$ | pJGXT01 -> SGB1003 |
| SGB1006 | W168 sacA::pJGlux03-P$_{veg}$-*luxABCDE-cm*$^{r}$ | pJGlux03 -> W168 |
| SGB1007 | W168 *sacA*::pAH328 | pAH328 -> W168 |
| SGB1011 | W168 *sacA*:: pJGlux01-P$_{merR20}$-*luxABCDE-cm*$^{r}$; *thrC*::pJGXT02- *P*$_{xylA}$-*MerRZntR- spc*$^{r}$ | pJGXT02 -> SGB1003 |
| SGB1014 | W168 *sacA*::pJGlux02- *P*$_{merR19}$-*luxABCDE-cm*$^{r}$ | pJGlux02 -> W168 |
| SGB1015 | W168 *sacA*:: pJGlux02- P$_{merR19}$-*luxABCDE-cm*$^{r}$; *thrC*:: pJGXT01- *P*$_{xylA}$-*MerR- spc*$^{r}$ | pJGXT02 ->SGB1014 |
| SGB1027 | W168 *sacA*:: pJGlux01-P$_{merR20}$-*luxABCDE-cm*$^{r}$ -*cm*$^{r}$; *thrC*:: pJGXT03- *P*$_{xylA}$-*MerRCueR -spc*$^{r}$ | pJGXT03 -> SGB1003 |
| SGB1028 | W168 *sacA*:: pJGlux01-P$_{merR}$-*luxABCDE-cm*$^{r}$; *thrC*:: pJGXT07- *P*$_{xylA}$-*MerRCueR*$^{A29T}$ *-spc*$^{r}$ | pJGXT07 -> SGB1003 |
| SGB1029 | W168 *sacA*:: pJGlux01-P$_{merR20}$-*luxABCDE-cm*$^{r}$; *thrC*:: pJGXT08- *P*$_{xylA}$-*MerRCueR*$^{A29T/G30P}$ *-spc*$^{r}$ | pJGXT08 -> SGB1003 |
| SGB1030 | W168 *sacA*:: pJGlux01-P$_{merR20}$-*luxABCDE-cm*$^{r}$; *thrC*:: pJGXT11-*P*$_{xylA}$-*MerRCueR*$^{A29T/G30P/P32M}$ *-spc*$^{r}$ | pJGXT11 -> SGB1003 |
| SGB1034 | W168 sacA::pJGlux04-P$_{cadA}$-*luxABCDE-cm*$^{r}$ | pJGlux04 -> W168 |
| SGB1035 | W168 *sacA*:: pJGlux01-P$_{merR20}$-*luxABCDE-cm*$^{r}$; *thrC*:: pJGXT15-P$_{xylA}$-MerRZntR$^{A29E/G30H}$ *-spc*$^{r}$ | pJGXT15 -> SGB1003 |
| SGB1036 | W168 *sacA*:: pJGlux01-P$_{merR20}$-*luxABCDE-cm*$^{r}$; *thrC*:: pJGXT16- *P*$_{xylA}$-*MerRZntR*$^{A29E/G30H}$ *-spc*$^{r}$ | pJGXT16 -> SGB1003 |
| SGB1047 | W168 *sacA*:: pJGlux01-P$_{merR20}$-*luxABCDE-cm*$^{r}$; *thrC*:: pJGXT23-P$_{xylA}$-MerRZntR$^{A29E/G30H/P32V}$ *-spc*$^{r}$ | pJGXT23 -> SGB1003 |
| SGB1050 | W168 *sacA*:: pJGlux02-P$_{merR19}$-*luxABCDE-cm*$^{r}$; *thrC*:: pJGXT11-P$_{xylA}$-MerRCueR$^{A29T/G30P/P32M}$ -*spc*$^{r}$ | pJGXT11 -> SGB1014 |
| SGB1060 | W168 *ΔoxdC-yvrJ-rsiO/sigO-rsoA* | pBSANDdel -> W168 |
| SGB1061 | W168 *ΔoxdC-yvrJ-rsiO/sigO-rsoA*; *sacA*::pBSANDlux-*cm*$^{r}$ | pBSANDlux -> SGB1060 |
| SGB1064 | W168 *ΔoxdC-yvrJ-rsiO/sigO-rsoA*; *sacA*::pBSANDlux-*cm*$^{r}$; *lacA*::pBSAND2-P$_{xylA}$-*MLS*$^{r}$ | pBSAND2-P$_{xylA}$ -> SGB1061 |
| SGB1065 | W168 *ΔoxdC-yvrJ-rsiO/sigO-rsoA*; *sacA*::pBSANDlux-*cm*$^{r}$; *lacA*::pBSAND2-P$_{xylA}$-*MLS*$^{r}$; *thrC*::pBSAND1-P$_{liaI}$-*spc*$^{r}$ | pBSAND1-P$_{liaI}$ -> SGB1064 |
| SGB1067 | W168 *sacA*::pBSGGlux-P$_{cadA}$-*cm*$^{r}$ | pBSGGlux-P$_{cadA}$ -> W168 |
| SGB1068 | W168 *ΔoxdC-yvrJ-rsiO/sigO-rsoA*; *sacA*::pBSANDlux -*cm*$^{r}$; *lacA*::pBSAND2-P$_{cadA}$-*MLS*$^{r}$ | pBSAND2-P$_{cadA}$ -> SGB1061 |
| SGB1072 | W168 *ΔoxdC-yvrJ-rsiO/sigO-rsoA*; *sacA*::pBSANDlux -*cm*$^{r}$; *lacA*::pBSAND2-P$_{cadA}$-*MLS*$^{r}$; *thrC*::pBSAND1-P$_{xylA}$-MerRZntR$^{A29E}$-Terminator-P$_{merR20}$-*spc*$^{r}$ | pBSAND1-P$_{xylA}$-MerRZntR$^{A29E}$-Terminator-P$_{merR}$ -> SGB1068 |

[a] Relevant characteristics are listed. Antibiotic resistance cassettes are denoted as follows: *cm*: chloramphenicol resistance; spc: spectinomycin resistance; MLS: erythromycin and lincomycin resistance.

[b] The direction of strain construction is indicated by an arrow which involves transformation with plasmids as indicated. Plasmids referred to are given in Table S2.

Supplementary Table S2. Plasmids used in this study

| Plasmid | Characteristics[a] | Source |
|---|---|---|
| pXT | Plasmid for xylose-inducible gene expression; integrates in *thrC*; spc$^r$, amp$^r$, ori ColE1 | [1] |
| pAH328 | Plasmid for transcriptional promoter fusions to *luxABCDE* (luciferase); integrates at *sacA*; cm$^r$, amp$^r$ | [2] |
| pBS4S | Empty plasmid, integration at *thrC*, amp$^r$, spec$^r$,, ori ColE1 | [3] |
| pBS2E | Empty plasmid, integration at *lacA*, amp$^r$, *mls*$^r$, ori ColE1 | [3] |
| pBS3Clux | Bacterial luciferase (*luxABCDE*) plasmid, integration at *sacA*, amp$^r$, cm$^r$, ori ColE1 | [3] |
| pSB1A3-$P_{xylA}$ | *amp*$^r$ , ori pMB1, $P_{xylA}$ (xylose inducible promoter) | [3] |
| pSB1C3-$P_{veg}$ | *cm*$^r$ *, ori pMB1,* $P_{veg}$ (strong constitutive B. subtilis promoter) | [3] |
| pJOE8999 | CRISPR-Cas9 deletion plasmid, *kan*$^r$; pUC-ori , temperature sensitive ori for *B. subtilis*(permissive at 30°C ) | [4] |
| pJGlux01 | $P_{merR20}$ – *luxABCDE* transcriptional fusion, derived from pAH328 | This study |
| pJGlux02 | $P_{merR19}$ – *luxABCDE* transcriptional fusion, derived from pAH328 | This study |
| pJGlux03 | $P_{veg}$-*luxABCDE* transcription fusion, derived from pAH328 | This study |
| pJGlux04 | $P_{cadA19}$-*luxABCDE* transcriptional fusion, derived from pAH328 | This study |
| pJGXT01 | $P_{xylA}$ -MerR transcriptional fusion, derived from pXT | This study |
| pJGUC01 | pUC19 derived vector, *amp*$^r$, ori ColE1, source of MerRZntR chimera RBS and CDS | This study (GenScript synthesised vector) |
| pJGXT02 | $P_{xylA}$ -MerRZntR transcriptional fusion, derived from pXT | This study |
| pJGUC02 | pUC19 derived vector, *amp*$^r$, ori ColE1, source of MerRCueR chimera RBS and CDS | This study (GenScript synthesised vector) |
| pJGXT03 | $P_{xylA}$ -MerRCueR transcriptional fusion, derived from pXT | This study |

| | | |
|---|---|---|
| pJGXT07 | $P_{xylA}$ -MerRCueR$^{A29T}$ transcriptional fusion, derived from pJGXT03 | This study |
| pJGXT08 | $P_{xylA}$ -MerRCueR$^{A29T/G30P}$ transcriptional fusion, derived from pJGXT03 | This study |
| pJGXT11 | $P_{xylA}$ -MerRCueR$^{A29T/G30P/P32M}$ transcriptional fusion, derived from pJGXT03 | This study |
| pJGXT15 | $P_{xylA}$ -MerRZntR$^{A29E}$ transcriptional fusion, derived from pJGXT02 | This study |
| pJGXT16 | $P_{xylA}$ -MerRZntR$^{A29E/G30H}$ transcriptional fusion, derived from pJGXT02 | This study |
| pJGXT23 | $P_{xylA}$ -MerRZntR$^{A29E/G30H/P32V}$ transcriptional fusion, derived from pJGXT02 | This study |
| pJGBS3Clux02 | Bacterial luciferase (*luxABCDE*) plasmid with BsaI deletion in *amp*$^{r}$, pBS3Clux derived | This study |
| pJGBS3Clux03 | Bacterial luciferase (*luxABCDE*) plasmid with BsaI deletion in *amp*$^{r}$ and *luxC*, derived from pJGBS3Clux02 | This study |
| pBSGGlux | Bacterial luciferase (*luxABCDE*) plasmid, integration at *sacA*, amp$^{r}$, cm$^{r}$, ori ColE1 – Golden Gate vector, derived from pJGBS3Clux03 | This study |
| pJGBS2E03 | Empty plasmid with BsaI deletion in *amp*$^{r}$ , derived from pBS2E | This study |
| pJGBS2E04 | Empty plasmid with BsaI deletion in *amp*$^{r}$ , and NgoMIV insertion in *amp*$^{r}$ derived from pBS2E | This study |
| pBSAND2 | Empty plasmid with Golden Gate cloning site allowing for SigO expression, derived from pJGBS2E04 | This study |
| pJGBS4S01 | Empty plasmid with BsaI deletion in *amp*$^{r}$ | This study |
| pBSAND1 | Empty plasmid with Golden Gate cloning site allowing for RsoA expression, derived from pJGBS4S01 | This study |
| pBSANDdel | CRISPR-Cas9 deletion vector allowing for guided cut upstream of *rsiO* in *B. subtilis* with repair homology flanking | This study |

| | | |
|---|---|---|
| | region to allow for *oxdC-yvrJ-rsiO/sigO-rsoA* deletion (SigO-RsoA regulon) | |
| pBSANDlux | $P_{oxdC}$ – luciferase reporter regulated by both SigO and RsoA, derived from pBSGGlux | This study |
| pBSAND2-$P_{xylA}$ | Xylose-inducible expression of SigO, derived from pBSAND2 | This study |
| pBSAND1-$P_{liaI}$ | Bacitracin-inducible expression of RsoA, derived from pBSAND1 | This study |
| pBSGGlux-$P_{cadA}$ | Metal-inducible expression of luciferase – natively regulated by CzrA in *B. subtilis* | This study |
| pBSAND2-$P_{cadA}$ | Metal-inducible expression of SigO, derived from pBSAND2 | This study |
| pBSAND1-$P_{xylA}$-MerRZntR$^{A29E}$-Terminator-$P_{merR20}$ | Xylose-inducible expression of MerRZntR$^{A29E}$ to subsequently regulate RsoA expression driven by $P_{merR20}$ (regulated by MerRZntR$^{A29E}$) – derived from pBSAND1 | This study |

[a] Relevant characteristics are listed. Antibiotic resistance cassettes as follows: *amp*$^r$ ; ampicillin resistance; *kan*$^r$ : kanamycin resistance; *cm*$^r$ : chloramphenicol resistance; spc$^r$ : spectinomycin resistance; *mls*: erythromycin and lincomycin resistance.

Supplementary Table S3. Primers used in this study

| Primer | Description | Sequence[a] | Function |
|---|---|---|---|
| SG0245 | pBS2E/pBSAND2 check fwd | GGCAACCGAGCGTTCTG | Verification of inserts into pBS2E derived plasmids |
| SG0246 | pBS2E/pBSAND2 check rev | CTGACAGCGTTTCGATCC | Verification of inserts into pBS2E derived plasmids |
| SG0528 | *sacA* insertion up fwd | CTGATTGGCATGGCGATTGC | Verification of *sacA* integration for pBS3Clux derived plasmids |
| SG0529 | *sacA* insertion up rev | ACAGCTCCAGATCCTCTACG | Verification of *sacA* integration for pBS3Clux derived plasmids |
| SG0530 | *sacA* insertion down fwd | GTCGCTACCATTACCAGTTG | Verification of *sacA* integration for pBS3Clux derived plasmids |
| SG0531 | *sacA* insertion down fwd | TCCAAACATTCCGGTGTTATC | Verification of *sacA* integration for pBS3Clux derived plasmids |
| SG0148 | pBS2E/pBSAND2 up fwd | GCATACCGGTTGCCGTCATC | Verification of lacA integration for pBS2E derived plasmids |
| SG1305 | pBS2E/pBSAND2 up rev | ATCTATTATTTAACGGGAGGAAATAATTC | Verification of lacA integration for pBS2E derived plasmids |
| SG1346 | pBS2E/pBSAND2 down fwd | CTGCAGAGATATCGATTTCAAGC | Verification of lacA integration for pBS2E derived plasmids |
| SG0149 | pBS2E/pBSAND2 down fwd | GAACTACATGCACTCCACAC | Verification of lacA integration for pBS2E derived plasmids |
| SG601 | pBS4S/pBSAND1 check fwd | CAGTCAACCCTTACCGCATTG | Verification of inserts into pBS4S derived plasmids |
| SG602 | pBS4S/pBSAND1 check fwd | CCTCCTCACTATTTTGATTAGTACC | Verification of inserts into pBS4S derived plasmids |
| SG0985 | EcoRI + $P_{merR}$ fwd | tttaa**GAATTC**AACGGAAGAATGTGGCTTCTTGG | Amplification of $P_{merR20}$ |
| SG0986 | SpeI + $P_{merR}$ rev | tttaa**ACTAGT**TGATTTTCATCCCCATATTGTCACC | Amplification of $P_{merR20}$ |
| SG0987 | BamHI + Native RBS + MerR fwd | tttaa**GGATCC**gaggtGACAATATGGGGATGAAAATC | Amplification of MerR with native RBS |
| SG0988 | EcoRI + MerR rev | tttaa**GAATTC**TTATTTATCAGGCCCTCCCATTAACGTT | Amplification of MerR with native RBS |
| SG0991 | pAH328/pBS3Clux check fwd | GAGCGTAGCGAAAAATCC | Verification of inserts into pAH328/pBS3Clux derived plasmids |
| SG0992 | pAH328/pBS3Clux check rev | GAAATGATGCTCCAGTAACC | Verification of inserts into pAH328/pBS3Clux derived plasmids |
| SG1124 | $P_{merR20}$ 1-bp deletion fwd | GGTACAGGGTTATACTTTTTATTGAGGTGACAATATGGGG | Mutagenesis of $P_{merR20}$ make $P_{merR19}$ |
| SG1125 | $P_{merR20}$ 1-bp deletion fwd | AAAAAGTATAACCCTGTACCATAGTACACGGTCAAGTCAT | Mutagenesis of $P_{merR20}$ make $P_{merR19}$ |

| | | | |
|---|---|---|---|
| SG1142 | MerRCueR$^{A29T}$ fwd | AGGATTGATAACAGGGCCTCCCAGAAACG AATCAGGGTATCGC | MerRCueR -> MerCueR$^{A29T}$ mutagenesis |
| SG1143 | MerRCueR$^{A29T}$ rev | TGGGAGGCCCTGTTATCAATCCTTTCCGCT CGTAATACCGAAC | MerRCueR -> MerCueR$^{A29T}$ mutagenesis |
| SG1154 | MerRCueR$^{A29T/G30P}$ fwd | ATTGATAACACCGCCTCCCAGAAACGAAT CAGGGTATCGCACC | MerRCueR -> MerRCueR$^{A29T/G30P}$ mutagenesis |
| SG1155 | MerRCueR$^{A29T/G30P}$ rev | TTCTGGGAGGCGGTGTTATCAATCCTTTCC GCTCGTAATACCG | MerRCueR -> MerRCueR$^{A29T/G30P}$ mutagenesis |
| SG1164 | P$_{cadA19}$ fwd | TTTAA**GAATTC**TTCGCGCTCGTAGTAGCG GATGGTC | Amplification of the P$_{cadA19}$ promoter from *P. aeruginosa* |
| SG1171 | P$_{cadA19}$ fwd | TTTAA**ACTAGT**TGTTGGGTGCGATTGCAC ACCCTGT | Amplification of the P$_{cadA19}$ promoter from *P. aeruginosa* |
| SG1167 | MerRCueR$^{A29T/G30P/P32M}$ fwd | AACACCGCCTATGAGAAACGAATCAGGGT ATCGCACCTACACG | MerRCueR$^{A29T/G30P}$ -> MerRCueR$^{A29T/G30P/P32M}$ mutagenesis |
| SG1168 | MerRCueR$^{A29T/G30P/P32M}$ rev | ATTCGTTTCTCATAGGCGGTGTTATCAATC CTTTCCGCTCGTA | MerRCueR$^{A29T/G30P}$ -> MerRCueR$^{A29T/G30P/P32M}$ mutagenesis |
| SG1172 | MerRZntR$^{A29E}$ fwd | AGGATTGATAGAAGGGCCTCCCAGAAAC GAATCAGGGTATCGA | MerRZntR -> MerRZntR$^{A29E}$ mutagenesis |
| SG1173 | MerRZntR$^{A29E}$ rev | TGGGAGGCCCTTCTATCAATCCTTTCCGCT CGTAATACCGAAC | MerRZntR -> MerRZntR$^{A29E}$ mutagenesis |
| SG1174 | MerRZntR$^{A29E/G30H}$ fwd | GAAAGGATTGATAGAACATCCTCCCAGAA ACGAATCAGGGTATCGACTA | MerRZntR -> MerRZntR$^{A29E/G30H}$ mutagenesis |
| SG1175 | MerRZntR$^{A29E/G30H}$ rev | CGTTTCTGGGAGG**ATGTTC**TATCAATCCTT TCCGCTCGTAATACCGAAC | MerRZntR -> MerRZntR$^{A29E/G30H}$ mutagenesis |
| SG1200 | MerRZntR$^{A29E/G30H/P32V}$ fwd | CGAGCGGAAAGGATTGATAGAACATCCTG TTAGAAACGAATCAGGGTATCGACTATAT ACC | MerRZntR -> MerRZntR$^{A29E/G30H/P32V}$ mutagenesis |
| SG1201 | MerRZntR$^{A29E/G30H/P32V}$ rev | CGATACCCTGATTCGTTTCTAACAGGATGT TCTATCAATCCTTTCCGCTCGTAATACC | MerRZntR -> MerRZntR$^{A29E/G30H/P32V}$ mutagenesis |
| SG1239 | BsaI deletion in *luxC* fwd | GACGGAATGAGGCCGTTGCAACGATTAGT GACATATATT | BsaI deletion in *luxC* of pBS3Clux |
| SG1240 | BsaI deletion in *luxC* rev | GTTGCAACGGCCTCATTCCGTCATGAGATC CACCAACTC | BsaI deletion in *luxC* of pBS3Clux |
| SG1242 | BsaI deletion in *amp$^r$* fwd | TGAGCGTGGCTCTCGCGGTATCATTGCAG CACTGG | BsaI deletion in *amp$^r$* |
| SG1243 | BsaI deletion in *amp$^r$* rev | ACCGCGAGAGCCACGCTCACCGGCTCCAG ATTTATCAG | BsaI deletion in *amp$^r$* |
| SG1245 | NgoMIV insertion in *amp$^r$* fwd | TTGACGCCGGCCAAGAGCAACTCGGTCGC CGCATAC | Insertion of NgoMIV site in *amp$^r$* |
| SG1246 | NgoMIV insertion in *amp$^r$* rev | GTTGCTCTTGGCCGGCGTCAACACGGGAT AATACC | Insertion of NgoMIV site in *amp$^r$* |
| SG1274 | SfiI + RBS + SigO fwd | TTAAA**GGCC**ACCCA**GGCC**gctcttaaggagga ttttagaATGAAGCATCCCATCGTGAAGCAT | Amplification of SigO to include RBS R1 |

| | | | |
|---|---|---|---|
| SG1248 | PstI + SigO + 3' - Terminator rev | AAA**CTGCAG***AAAAAACGCGAGCGTCAGGCGCTCGCGTTTTT*CTAATCGATTACTTTTCCCCCTTTTTTTGCGC | Amplification of SigO to include 3'-Terminator |
| SG1275 | SfiI + RBS + RsoA fwd | TTAAA**GGCC**ACCCA**GGCC**gctcttaaggaggattttagaGTGGACGGCCAGTTTGAACAAAA | Amplification of RsoA to include RBS R1 |
| SG1250 | PstI + RsoA + 3'- Terminator rev | TTAAA**CTGCAG**AAAAAACGCGAGCGTCAGGCGCTCGCGTTTTTCTAATCGATTATGAGTTCTCATCTACCATGTAC | Amplification of RsoA to include 3' - Terminator |
| SG1272 | EcoRI + RFP + 5'- Terminator + 5'-BsaI fwd | TTAAA**GAATTC**AAAAACGCGAGCGCCTGACGCTCGCGTTTTTTCC**C**TACTC**GAGACC**TCATTCTCGCAATACGCAAACCGCCTCTC | Amplification of RFP to include 5'-Terminator and 5' BsaI site (new BsaI Golden Gate site) – For plasmids pBSAND1/2 and pBSANDlux |
| SG1273 | SfiI + RFP + 5'-BsaI site rev | TTAAA**GGCC**TGGGT**GGCC**TACTGATAATTCT**GAGACC**ATAAACGCAGAAAGGCCCACCC | Amplification of RFP to include 3'-BsaI site (new BsaI Golden Gate site) – For plasmids pBSAND1/2 |
| SG1324 | PstI + RFP + 3'- BsaI site rev | TTAAA**CTGCAG**GGTACTGATAATTCT**GAGACC**ATAAACGCAGAAAGGCCCACCC | Amplification of RFP to include 3'-BsaI site (new Golden Gate site) – for plasmid pBSANDlux |
| SG1297 | BsaI + $P_{xylA}$ fwd | TTTAAA**GGTCTC**ATACTAAGGCCAAAAAACTGCTGCCTTCG | Amplification of $P_{xylA}$ |
| SG1298 | BsaI + $P_{xylA}$ rev | TTTAAA**GGTCTC**TATTCTTCGATAAGCTTGGGATCCCAG | Amplification of $P_{xylA}$ |
| SG1299 | BsaI + $P_{oxdC}$ fwd | TTTAAA**GGTCTC**ATACTGATTCGAAAAGAAGTTTGATCA | Amplification of $P_{oxdC}$ |
| SG1300 | BsaI + $P_{oxdC}$ rev | TTTAAA**GGTCTC**TATTCTTCGTTTTCCCTTCTGTTTTGT | Amplification of $P_{oxdC}$ |
| SG1303 | pBSGGlux new GoldenGate site check fwd | AAAAACGCGAGCGCCTGAC | Check for insertion of RFP with new Golden Gate site in pBSGGlux |
| SG1325 | pBSGGlux new GoldenGate site check rev | GCAGGGTACTGATAATTCTGAGACCAT | Check for insertion of RFP with new Golden Gate site in pBSGGlux |
| SG1326 | *oxdC-yvrJ-rsiO/sigO-rsoA* left hand homology + SfiI fwd | Ttaaa**GGCC**AACGA**GGCC**TCAATGAAGCTGTCAGAGATGAAC | Amplification of *oxdC-yvrJ-rsiO/sigO-rsoA*left hand homology region for pBSANDdel deletion vector |
| SG1327 | *oxdC-yvrJ-rsiO/sigO-rsoA* left hand homology + SfiI rev | TTAAA**GGCC**TAAAT**GGCC**AAGAACGCATTCCAATTGGATGC | Amplification of *oxdC-yvrJ-rsiO/sigO-rsoA* left hand homology region for pBSANDdel deletion vector |
| SG1328 | *oxdC-yvrJ-rsiO/sigO-rsoA* right hand homology + SfiI fwd | TTAAA**GGCC**ATTTA**GGCC**TTTCATATGAAAAAACATCAAAAGGGAG | Amplification of *oxdC-yvrJ-rsiO/sigO-rsoA* right hand homology region for pBSANDdel deletion vector |

| | | | |
|---|---|---|---|
| SG1329 | *oxdC-yvrJ-rsiO/sigO-rsoA* right hand homology + SfiI rev | TTAAA**GGCC**TTATT**GGCC**GCCTACGATCA ACTCGGCATT | Amplification of *oxdC-yvrJ-rsiO/sigO-rsoA* right hand homology region for pBSANDdel deletion vector |
| SG1344 | *ΔoxdC-yvrJ-rsiO/sigO-rsoA* deletion check fwd | GAACATATTCGAATCTCCTTTCAA | Check for *ΔoxdC-yvrJ-rsiO/sigO-rsoA* deletion using pBSANDdel |
| SG1345 | *ΔoxdC-yvrJ-rsiO/sigO-rsoA* deletion check rev | TGCCTTCAGGATTTGGGATAAA | Check for *ΔoxdC-yvrJ-rsiO/sigO-rsoA* deletion using pBSANDdel |
| SG1347 | pBSANDdel homology insertion check fwd | GATTTGTTCAGAACGCTCGGT | Check for insertion of *oxdC-yvrJ-rsiO/sigO-rsoA* flanking homology regions into pBSANDdel |
| SG1348 | pBSANDdel homology insertion check rev | ACGCATTGATTTGAGTCAGCT | Check for insertion of *oxdC-yvrJ-rsiO/sigO-rsoA* flanking homology regions into pBSANDdel |
| SG1349 | sgRNA for *ΔoxdC-yvrJ-rsiO/sigO-rsoA* deletion fwd | TACGGTTCTCTGCAAGCCTTCATA | sgRNA to guide Cas9 in pBSANDdel for *ΔoxdC-yvrJ-rsiO/sigO-rsoA* deletion |
| SG1350 | sgRNA for *ΔoxdC-yvrJ-rsiO/sigO-rsoA* deletion fwd | AAACTATGAAGGCTTGCAGAGAAC | sgRNA to guide Cas9 in pBSANDdel for *ΔoxdC-yvrJ-rsiO/sigO-rsoA* deletion |
| SG1382 | MerRZntR$^{A29E}$ + BsaI fwd | TTTAAA**GGTCTC**GTTTTgaggtGACAATATG GGGATGAAAA | Amplification of MerRZntR$^{A29E}$ for Golden Gate assembly |
| SG1383 | MerRZntR$^{A29E}$ + BsaI rev | TTTAAA**GGTCTC**AGGGGTCAACAACCACT CTTAACGCCACT | Amplification of MerRZntR$^{A29E}$ for Golden Gate assembly |
| SG1384 | $P_{merR}$ with 5'-Terminator + BsaI fwd | TTTAAA**GGTCTC**ACCCCAAAAACGCGAGC GCCTGACGCTCGCGTTTTTTAACGGAAGA ATGTGGCTTCTTGG | Amplification of $P_{merR}$ to include 5'-Terminator |
| SG1385 | $P_{merR}$ with 5'-Terminator + BsaI rev | TTTAAA**GGTCTC**TATTCTGATTTTCATCCCC ATATTGTCACC | Amplification of $P_{merR}$ to include 5'-Terminator |
| SG1387 | $P_{cadA}$ + BsaI Rev | TTTAAA**GGTCTC**ATACTGCGCTTGCTGTTT TTCATTGACACT | Amplification of $P_{cadA}$ |
| SG1403 | $P_{cadA}$ + BsaI Fwd | TTTAAA**GGTCTC**ATACTATTCAGCTCCGTT TCCGTTGTTCT | Amplification of $P_{cadA}$ |
| SG1388 | $P_{liaI}$ + BsaI fwd | TTTAAA**GGTCTC**ATACT*ATTGGCCAAAGCA GAAAGGTC* | Amplification of $P_{liaI}$ |
| SG1389 | $P_{liaI}$ + BsaI rev | TTTAAA**GGTCTC**ATACT*ATTGGCCAAAGCA GAAAGGTC* | Amplification of $P_{liaI}$ |
| SG1410 | $P_{xylA}$ + BsaI rev | TTTAAA**GGTCTC**CAAAATTCGATAAGCTTG GGATCCCAG | Amplification of $P_{xylA}$ |

[a] Restriction sites are in uppercase bold; QuickChange point mutation sites are underlined bold, RBS sequences in primers are in lowercase and underlined with a thin line; terminator sequence are underlined with a dotted line.

Supplementary Table S4. Genetic Sequences used in this study

| Part | Part type & Function | Sequence[a] |
|---|---|---|
| $P_{merR20}$ | MerR Family promoter recognised by MerR and the chimeras derived from this study. Contains a 20-bp spacer between the -10 and -35 elements | AACGGAAGAATGTGGCTTCTTGGCGTGAAGAGCAGA<br>TATCTTTATTCTTAAACTTCTAAAAAAGCTTATGTGAA<br>CACTTGATAAATAAGGTTTTTTATCTGACCTATTTTAT<br>AAGATTATTCTATAAAAGAAAAAATAATGTATGAC**TT**<br>**GACC**GTGTACTATGGTACAGGTT**TATACT**TTTTATT<br>GAGGTGACAATATGGGGATGAAAATCAGTGAATTGG<br>CTAAAGCGTGTGATGTGAATAAAGA |
| $P_{merR19}$ | MerR Family promoter recognised by MerR and the chimeras derived from this study. Contains a 19-bp spacer between the -10 and -35 elements | AACGGAAGAATGTGGCTTCTTGGCGTGAAGAGCAGA<br>TATCTTTATTCTTAAACTTCTAAAAAAGCTTATGTGAA<br>CACTTGATAAATAAGGTTTTTTATCTGACCTATTTTAT<br>AAGATTATTCTATAAAAGAAAAAATAATGTATGAC**TT**<br>**GACC**GTGTACTATGGTACAGGGT**TATACT**TTTTATTG<br>AGGTGACAATATGGGGATGAAAATCAGTGAATTGGC<br>TAAAGCGTGTGATGTGAATAAAGA |
| $P_{veg}$ | Strong constitutive promoter native to B. subtilis | GGAGTTCTGAGAATTGGTATGCCTTATAAGTCCAATT<br>AACAGTTGAAAACCTGCATAGGAGAGCTATGCGGGT<br>TTTTTATTTTACATAATGATACATAATTTACCGAAACTT<br>GCGGAACATAATTGAGGAATCATAGAATTTTGTCAAA<br>ATAATTTTATTGACAACGTCTTATTAACGTTGATATAA<br>TTTAAATTTTAT**TTGACA**AAAATGGGCTCGTGTTG**TA**<br>**CAAT**AAATGTAGT |
| MerR | Coding sequence for MerR including the native RBS and spacer region of MerR. Hg$^{2}$-sensor | gaggtgacaatATGGGGATGAAAATCAGTGAATTGGCT<br>AAAGCGTGTGATGTGAATAAAGAAACCGTTCGGTAT<br>TACGAGCGGAAAGGATTGATAGCCGGGCCTCCCAGA<br>AACGAATCAGGGTATCGAATATATTCAGAGGAAACA<br>GCAGATCGGGTACGGTTTATTAAACGAATGAAGGAA<br>TTGGATTTCTCGCTAAAGGAAATCCACCTGTTGTTTG<br>GTGTGGTTGATCAAGATGGGGAGAGATGTAAAGATA<br>TGTACGCCTTTACCGTTCAAAAAACCAAAGAAATCGA<br>GCGGAAAGTGCAGGGTTTGTTACGAATCCAACGGTT<br>ATTAGAGGAATTAAAAGAAAAGTGTCCAGATGAAAA<br>GGCGATGTATACCTGTCCTATTATTGAAACGTTAATG<br>GGAGGGCCTGATAAATAA |
| MerRZntR | Coding sequence for MerRZntR chimera including the native RBS and spacer region of MerR | gaggtgacaatATGGGGATGAAAATCAGTGAATTGGCT<br>AAAGCGTGTGATGTGAATAAAGAAACCGTTCGGTAT<br>TACGAGCGGAAAGGATTGATAGCCGGGCCTCCCAGA<br>AACGAATCAGGGTATCGACTATATACCGAAAGCGAT<br>CTCCAGCGATTGAAATTTATCCGCCATGCCAGACAAC<br>TAGGTTTCAGTCTGGAGTCGATCCGCGAGTTGCTGTC<br>GATCCGCATCGATCCTGAACACCATACCTGTCAGGAG<br>TCAAAAGGCATTGTGCAGGAAAGATTGCAGGAAGTC<br>GAAGCACGGATAGCCGAGTTGCAGAGTATGCAGCGT<br>TCCTTGCAACGCCTTAACGATGCCTGTTGTGGGACTG<br>CTCATAGCAGTGTTTATTGTTCGATTCTTGAAGCTCTT<br>GAACAAGGGGCGAGTGGCGTTAAGAGTGGTTGTTGA |

| | | |
|---|---|---|
| MerRZntR$^{A29E}$ | Coding sequence for MerRZntR chimera including the native RBS and spacer region of MerR. Contains a A29E substitution in the α-helix 2-3 loop region. Optimised Zn$^{2+}$ - sensor | gaggtgacaatATGGGGATGAAAATCAGTGAATTGGCT<br>AAAGCGTGTGATGTGAATAAAGAAACCGTTCGGTAT<br>TACGAGCGGAAAGGATTGATAGAAGGGCCTCCCAGA<br>AACGAATCAGGGTATCGACTATATACCGAAAGCGAT<br>CTCCAGCGATTGAAATTTATCCGCCATGCCAGACAAC<br>TAGGTTTCAGTCTGGAGTCGATCCGCGAGTTGCTGTC<br>GATCCGCATCGATCCTGAACACCATACCTGTCAGGAG<br>TCAAAAGGCATTGTGCAGGAAAGATTGCAGGAAGTC<br>GAAGCACGGATAGCCGAGTTGCAGAGTATGCAGCGT<br>TCCTTGCAACGCCTTAACGATGCCTGTTGTGGGACTG<br>CTCATAGCAGTGTTTATTGTTCGATTCTTGAAGCTCTT<br>GAACAAGGGGCGAGTGGCGTTAAGAGTGGTTGTTGA |
| MerRZntR$^{A29E/G30H}$ | Coding sequence for MerRZntR chimera including the native RBS and spacer region of MerR. Contains a A29E/G30H substitution in the α-helix 2-3 loop region | gaggtgacaatATGGGGATGAAAATCAGTGAATTGGCT<br>AAAGCGTGTGATGTGAATAAAGAAACCGTTCGGTAT<br>TACGAGCGGAAAGGATTGATAGAACATCCTCCCAGA<br>AACGAATCAGGGTATCGACTATATACCGAAAGCGAT<br>CTCCAGCGATTGAAATTTATCCGCCATGCCAGACAAC<br>TAGGTTTCAGTCTGGAGTCGATCCGCGAGTTGCTGTC<br>GATCCGCATCGATCCTGAACACCATACCTGTCAGGAG<br>TCAAAAGGCATTGTGCAGGAAAGATTGCAGGAAGTC<br>GAAGCACGGATAGCCGAGTTGCAGAGTATGCAGCGT<br>TCCTTGCAACGCCTTAACGATGCCTGTTGTGGGACTG<br>CTCATAGCAGTGTTTATTGTTCGATTCTTGAAGCTCTT<br>GAACAAGGGGCGAGTGGCGTTAAGAGTGGTTGTTGA |
| MerRZntR$^{A29E/G30H/P32V}$ | Coding sequence for MerRZntR chimera including the native RBS and spacer region of MerR. Contains a A29E/G30H/P32V substitution in the α-helix 2-3 loop region | gaggtgacaatATGGGGATGAAAATCAGTGAATTGGCT<br>AAAGCGTGTGATGTGAATAAAGAAACCGTTCGGTAT<br>TACGAGCGGAAAGGATTGATAGAACATCCTGTTAGA<br>AACGAATCAGGGTATCGACTATATACCGAAAGCGAT<br>CTCCAGCGATTGAAATTTATCCGCCATGCCAGACAAC<br>TAGGTTTCAGTCTGGAGTCGATCCGCGAGTTGCTGTC<br>GATCCGCATCGATCCTGAACACCATACCTGTCAGGAG<br>TCAAAAGGCATTGTGCAGGAAAGATTGCAGGAAGTC<br>GAAGCACGGATAGCCGAGTTGCAGAGTATGCAGCGT<br>TCCTTGCAACGCCTTAACGATGCCTGTTGTGGGACTG<br>CTCATAGCAGTGTTTATTGTTCGATTCTTGAAGCTCTT<br>GAACAAGGGGCGAGTGGCGTTAAGAGTGGTTGTTGA |
| MerRCueR | Coding sequence for MerRCueR chimera including the native RBS and spacer region of MerR | gaggtgacaatATGGGGATGAAAATCAGTGAATTGGCT<br>AAAGCGTGTGATGTGAATAAAGAAACCGTTCGGTAT<br>TACGAGCGGAAAGGATTGATAGCCGGGCCTCCCAGA<br>AACGAATCAGGGTATCGCACCTACACGCAGCAGCAT<br>CTCAACGAACTGACCTTACTGCGCCAGGCACGGCAG<br>GTGGGCTTTAACCTGGAAGAGAGCGGCGAGCTGGTG<br>AATCTGTTTAACGACCCGCAGCGGCACAGCGCCGAC<br>GTCAAACGGCGCACGCTGGAGAAGGTGGCGGAGAT<br>CGAACGACACATTGAGGAGCTGCAATCCATGCGCGA<br>CCAGCTGCTGGCACTGGCGAATGCCTGCCCTGGCGA<br>TGACAGCGCCGACTGCCCGATTATCGAAAATCTCTCC<br>GGCTGCTGTCATCATCGGGCAGGGTGA |

| | | |
|---|---|---|
| MerRCueR$^{A29T}$ | Coding sequence for MerRCueR chimera including the native RBS and spacer region of MerR. Contains a A29T substitution in the α-helix 2-3 loop region | gaggtgacaatATGGGGATGAAAATCAGTGAATTGGCT<br>AAAGCGTGTGATGTGAATAAAGAAACCGTTCGGTAT<br>TACGAGCGGAAAGGATTGATAACAGGGCCTCCCAGA<br>AACGAATCAGGGTATCGCACCTACACGCAGCAGCAT<br>CTCAACGAACTGACCTTACTGCGCCAGGCACGGCAG<br>GTGGGCTTTAACCTGGAAGAGAGCGGCGAGCTGGTG<br>AATCTGTTTAACGACCCGCAGCGGCACAGCGCCGAC<br>GTCAAACGGCGCACGCTGGAGAAGGTGGCGGAGAT<br>CGAACGACACATTGAGGAGCTGCAATCCATGCGCGA<br>CCAGCTGCTGGCACTGGCGAATGCCTGCCCTGGCGA<br>TGACAGCGCCGACTGCCCGATTATCGAAAATCTCTCC<br>GGCTGCTGTCATCATCGGGCAGGGTGA |
| MerRCueR$^{A29T/G30P}$ | Coding sequence for MerRCueR chimera including the native RBS and spacer region of MerR. Contains a A29T/G30P substitution in the α-helix 2-3 loop region | gaggtgacaatATGGGGATGAAAATCAGTGAATTGGCT<br>AAAGCGTGTGATGTGAATAAAGAAACCGTTCGGTAT<br>TACGAGCGGAAAGGATTGATAACACCGCCTCCCAGA<br>AACGAATCAGGGTATCGCACCTACACGCAGCAGCAT<br>CTCAACGAACTGACCTTACTGCGCCAGGCACGGCAG<br>GTGGGCTTTAACCTGGAAGAGAGCGGCGAGCTGGTG<br>AATCTGTTTAACGACCCGCAGCGGCACAGCGCCGAC<br>GTCAAACGGCGCACGCTGGAGAAGGTGGCGGAGAT<br>CGAACGACACATTGAGGAGCTGCAATCCATGCGCGA<br>CCAGCTGCTGGCACTGGCGAATGCCTGCCCTGGCGA<br>TGACAGCGCCGACTGCCCGATTATCGAAAATCTCTCC<br>GGCTGCTGTCATCATCGGGCAGGGTGA |
| MerRCueR$^{A29T/G30P/P32M}$ | Coding sequence for MerRCueR chimera including the native RBS and spacer region of MerR. Contains a A29T/G30P/P32M substitution in the α-helix 2-3 loop region | gaggtgacaatATGGGGATGAAAATCAGTGAATTGGCT<br>AAAGCGTGTGATGTGAATAAAGAAACCGTTCGGTAT<br>TACGAGCGGAAAGGATTGATAACACCGCCTATGAGA<br>AACGAATCAGGGTATCGCACCTACACGCAGCAGCAT<br>CTCAACGAACTGACCTTACTGCGCCAGGCACGGCAG<br>GTGGGCTTTAACCTGGAAGAGAGCGGCGAGCTGGTG<br>AATCTGTTTAACGACCCGCAGCGGCACAGCGCCGAC<br>GTCAAACGGCGCACGCTGGAGAAGGTGGCGGAGAT<br>CGAACGACACATTGAGGAGCTGCAATCCATGCGCGA<br>CCAGCTGCTGGCACTGGCGAATGCCTGCCCTGGCGA<br>TGACAGCGCCGACTGCCCGATTATCGAAAATCTCTCC<br>GGCTGCTGTCATCATCGGGCAGGGTGA |
| RsoA | Coding sequence for RsoA, co-σ-factor of SigO | GTGGACGGCCAGTTTGAACAAAAAAAGAAACAAAAA<br>GACGAGACTTATGACATTGAGCACCTGATTGCATGCT<br>TTTCACCGATGATCAGA<br>AAAAAACTCAGCAATACGTCCTATCAAGAAAGAGAA<br>GATTTAGAGCAAGAGCTGAAGATCAAAATGTTTGAA<br>AAGGCTGATATGCTTTTATGTCAGGATGTACCGGGGT<br>TTTGGGAGTTTATTTTGTACATGGTAGATGAGAACTC<br>ATAA |
| SigO | Coding sequence for SigO, co-σ-factor of RsoA | ATGAAGCATCCCATCGTGAAGCATTTTTTGAGCAATC<br>CTCAGCATTACCGTTTGTTCAAAAACGTAATGGAAAG<br>CCCTAACGAAAAAGATGCAAGATCATTGGACGAGCT<br>ATTTAAGCAATTTTATAAAGAAATCCGCATCGTCAAG |

| | | |
|---|---|---|
| | | TATATGAATTCAATGATTCGCATCTTTTCTATTGATTTTGATAAGCGGGTTCGCAAAAACCAAAAACGGTATCCACTGACGGTTGATCATCCGGAGGCGGGAGATCGGCTTTCTTCCGAAACAGGTAGCGATGCATTTGAAGAATTTTTAGACAGGCAGGATGATTTGAGCCAGCATGTACAGGATTACCAGCTCTACCAAGCGATCCAGAAGCTGACTGACAAACAAAAAAGTGTGCTGACGAAAGTCTATCTTCACGGTGCCACGATGCAGGAGATTGCAGATTCATTAGGGGAGTCCCGACAAAACATCTCCAACATTCATAAAAAGGGGCTGGAGAATATCAGAAAGCAGTTAGCGGCGCAAAAAAAGGGGGAAAAGTAA |
| $P_{oxdC}$ | $P_{oxdC}$ promoter recognised by SigO and RsoA | GATTCGAAAAGAAGTTTGATCAACTAATAGAACTAATGACAGAACTGAAAGATCATGCAAAAAAATAATTTTTCAATCGAAG**TTGACT**TTTCACTGGTTTTTT**TCACTTA**ACAAAACAGAAGGGAAAACGAA |
| $P_{xylA}$ | Xylose-inducible promoter lacking CRE-element. Sequence is derived from Radeck *et al* [3] | AAGGCCAAAAAACTGCTGCCTTCGGATCAGCGATATCCACTTCATCCACTCCATTTGTTTAATCTTTAAATTAAGTATCAACATAGTACATAGCGAATCTTCCCTTTATTATATCTAATGTGTTCATAAAAAACTAAAAAAAA**TATTGA**AAATACTGACGAGGTTA**TATAAG**ATGAAAATAAGTTAGTTTGTTTAAACAACAAACTAATAGGTGATGTACTTACTATATGAAATAAAATGCATCTGGGATCCCAAGCTTATCGAA |
| $P_{liaI}$ | Bacitracin-inducible promoter. Sequence is derived from Radeck *et al* [3] | ATTGGCCAAAGCAGAAAGGTCCGACCTAATTAAAGAAAGGGAAGCAAGTGTTCATCTGTAAAGGGTTTTAAAACGCCATGCCTCGTGCATGGCGTTTTTTTGTGCCAATGGGTCCGGTGCGAGATACGACTCCGGTCTTATATAAAAATCAATCTCTGAT**TCG**TTTTGCATATCTTCCAACTTG**TATAAG**ATGAAGACAAGGAAAACGA |
| Terminator 1 “Term 1” | Strong minimal terminator to prevent transcriptional read-through. Sequence is derived from Cui *et al* [5] | AAAAACGCGAGCGCCTGACGCTCGCGTTTTTT |
| RBS R1 | RBS used for SigO and RsoA translation. Sequence is derived from Guiziou *et al* [6] | GCTCTTAAAGGGGGTTTTAGA |
| $P_{cadA}$ | Metal inducible promoter regulated natively by CzrA | ATTCAGCTCCGTTTCCGTTGTTCTGAATGCTCTTCGTCTGCAAAAAGTAAAATGAAAAACCGGCTATATGCCGGTTTTTGTTTTTCA**TTGACA**CTTTCTTGGAAAACAACA**TATAAT**AGGTGTAACTTATATATGAGTATATGCTCATATATATAAAATAAATACAATACTCATTGATACGCTTTGAAGAGGGAA |

[a] For sequences of this study, the -10 and -35 elements are in bold; positions where a regulator binds are underlined; RBS and spacer are in lowercase.

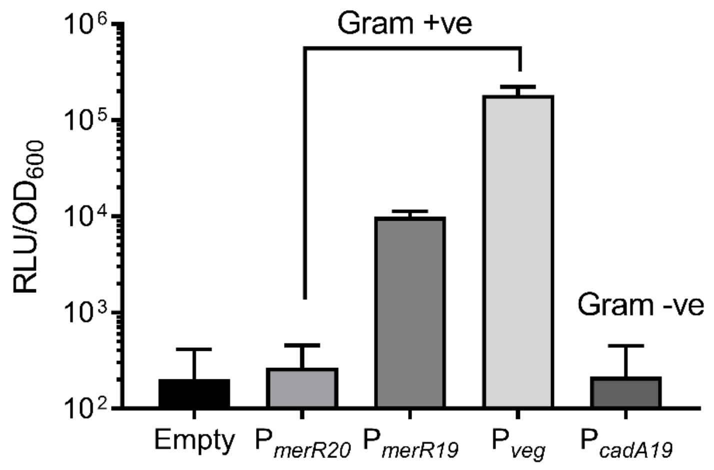


**Supplementary Figure S1. Comparison of Gram +ve and Gram -ve MerR promoter activity in *B. subtilis*.** Cells harbouring either an empty luciferase reporter, $P_{merR20}$ (Gram +ve), $P_{merR19}$ (Gram +ve), $P_{veg}$ (Gram +ve) or $P_{cadA19}$ (Gram-ve) were grown in MM9 media with luciferase activity measured overtime. Values presented are the average of three time points (35-, 40- and 45-minutes) following initial inoculation into MM9 medium. The strong Gram +ve *B. subtilis* promoter $P_{veg}$, was included as a control. Subscripts indicate the size of the spacer region between the -10 and -35 elements for MerR family promoters. Data are the ± standard deviation of triplicate measurements performed on three different days.

**A)**

```
ZntR_Ec    MYRIGELAKMAEVTPDTIRYYEKQQMMEHEVRTEGGFRLYTESDLQRLKFIRHARQLGFS 60
ZntR_Se    MYRIGELAKMAEVTPDTIRYYEKQQMMEHEVRTEGGFRLYTESDLQRLKFIRHARQLGFS 60
ZntR_Kp    MYRIGELAKMAEVTPDTIRYYEKQQMMEHEVRTEGGFRLYTESDLQRLKFIRHARQLGFS 60
ZntR_Sf    MYRIGELAKMAEVTPDTIRYYEKQQMMEHEVRTEGGFRLYTESDLQRLKFIRHARQLGFS 60
ZntR_Pa    MYRIGELAKLANVTPDTIRYYEKQQMIDHEVRTEGGFRLYTDNDLQRLRFIRYARQLGFT 60
           *********:*:**************::*************:.*****:***:******:
```

**B)**

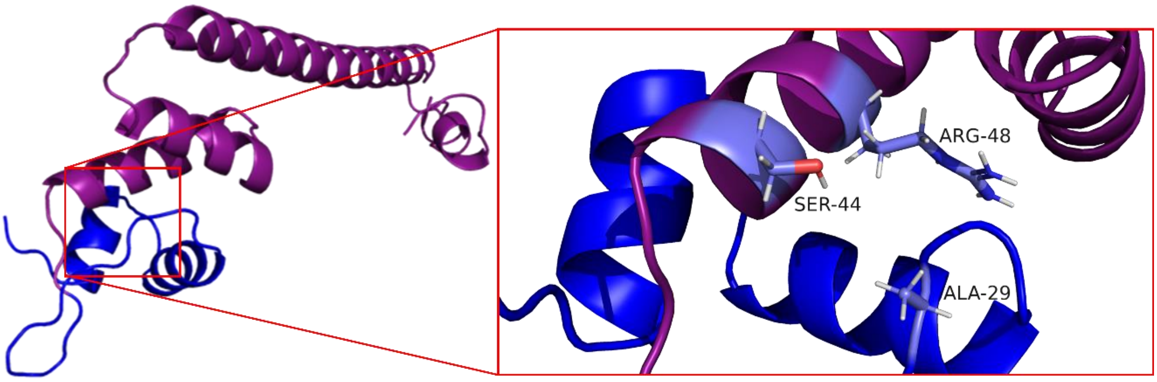


**Supplementary Figure S2. Comparison of ZntR sequences and structural analysis of the chimera MerRZntR. A)** Sequence alignment of ZntR homologues from various Gram-negative genetic backgrounds ("Ec" – *E. coli*, accession code: AAC76317.1; "Se"- *S. enterica*, accession code: EBW6030787.1; "Kp" – *K. pneumoniae*, accession code: CDK70471.1; "Sf" – *S. flexerni*, accession code: EAA3112577.1; "Pa" – *P. aeruginosa*, accession code: MBH4409345.1). Residues of interest involved in interdomain communication are highlighted in bold purple. Asterisk " * " indicate fully conserved residues, colon " : " indicates conserved residues with similar properties, and period " . " indicates residues of weakly similar properties. **B)** Structural analysis of the residues between α-helices 2-3 in MerRZntR. Here, the MerR (*S. aureus*) derived DNA-Binding Domain is coloured dark blue, whilst the ZntR (*E. coli*) derived Metal-Binding Domain is coloured purple. Residues of interest are coloured lavender blue and are numbered accordingly. Due to the presence of the non-polar residue Ala-29, no hydrogen bonding is present between Ala-29 (DNA-Binding Domain) and Ser-44 and Arg-48 (Metal Binding Domain).

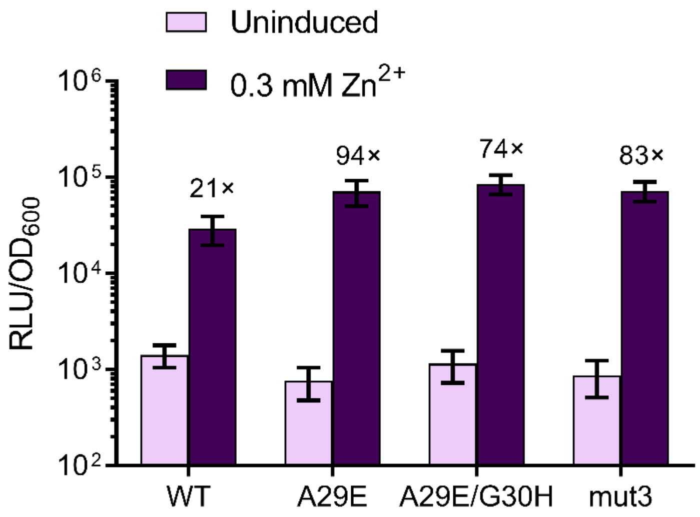


**Supplementary Figure S3. Activity of double and triple MerRZntR mutants against the wild-type and single mutant MerRZntR$^{A29E}$**. Residues in between alpha-helix 2-3 were mutated to those found natively in ZntR, generating MerRZntR$^{A29E/G30H}$ and MerRZntR$^{A29E/G30H/P32V}$ (mut3), with the activity compared relative to both wild-type (MerRZntR) and the single mutant (MerRZntR$^{A29E}$). Cells were grown to $OD_{600}$= ~ 0.03 and induced at the highest sub-lethal tested concentration of $Zn^{2+}$ with luciferase activity (relative luminescence units (RLU) normalised by cell density ($OD_{600}$)) for three time points (35-, 40- and 45-mins) post induction. Fold-induction values of the induced (dark purple) are relative to the respective uninduced strain (light purple). Values are presented as mean and ± standard deviation of either two or three independent replicates.

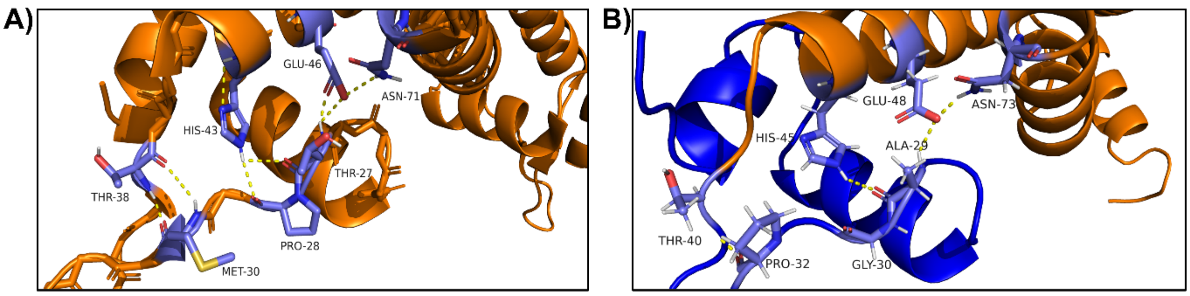


C)
```
CueR_Ec     MNISDVAKITGLTSKAIRFYEEKGLVTPPMRSENGYRTYTQQHLNELTLLRQARQVGFNL 60
CueR_Kp     MNISDVAKITGLTSKAIRFYEEKGLVTPPMRSENGYRTYTQQHLNELTLLRQARQVGFNL 60
CueR_Sf     MNISDVAKITGLTSKAIRFYEEKGLVTPPMRSENGYRTYTQQHLNELTLLRQARQVGFNL 60
CueR_Pa     MNISDVAKKTGLTSKAIRFYEEKGLVTPPLRSENGYRTYSQQHLDELTLLRQARQVGFNL 60
CueR_Se     MNISDVAKKTGLTSKAIRFYEEKGLVTPPLRSENGYRTYTQKHLNELTLLRQARQVGFNL 60
            ******** ********************:*********:*:**:***************
```

**Supplementary Figure S4. Comparison of inter-domain communication between CueR and the chimera MerRCueR. A)** Magnification of the CueR (*E. coli*, accession code: CAD6020341.1) inter-domain hydrogen bonding network between α-helices 2-3. **B)** Magnification of the MerRCueR chimera inter-domain hydrogen bonding network between α-helices 2-3. Here, the MerR (*S. aureus*) derived DNA-Binding Domain is indicated in dark blue, whilst the CueR (*E. coli*) derived Metal-Binding Domain is indicated in orange. For panels **A** and **B,** residues of interest colour in lavender blue and are numbered accordingly, with hydrogen bonds indicated in yellow. **C)** Sequence alignment of CueR from various Gram-negative genetic backgrounds ("Ec" – *E. coli*, accession code: CAD6020341.1; ; "Kp" – K. pneumoniae, accession code: OZQ58601.1; "Sf" – *S. flexerni*, accession code: EFX2973845.1; "Pa" – *P. aeruginosa*, accession code: MXH36715.1; "Se"- *S. enterica*, accession code: EAS1883030.1). Residues of interest involved in interdomain communication are highlighted in bold orange. Asterisk " * " indicate fully conserved residues, colon " : " indicates conserved residues with similar properties, and period " . " indicates residues of weakly similar properties.

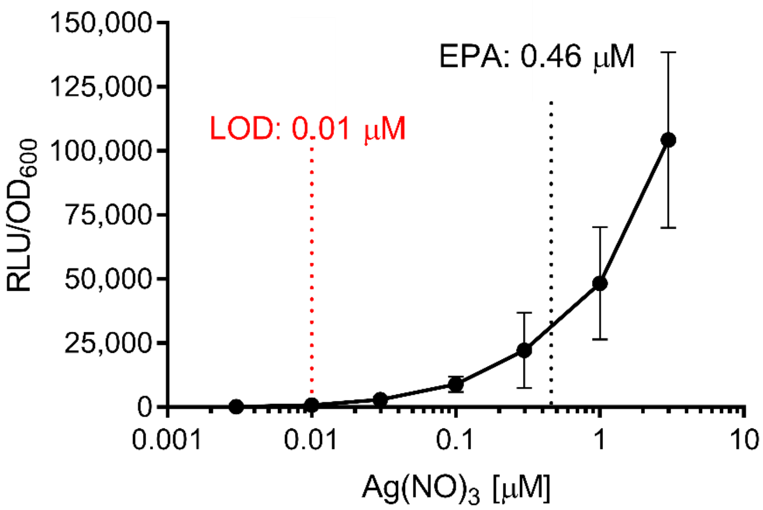


**Supplementary Figure S5. Dose response of $P_{merR}$ regulated by $MerRCueR^{mut3}$ in response to $Ag^{+}$ induction**. Transcriptional output from $P_{merR}$ is shown in response to various concentrations of $Ag^{+}$. Cells were induced at $OD_{600}$ = ~ 0.03 with luciferase activity (relative luminescence units [RLU]) normalised to optical density ($OD_{600}$) values ($RLU/OD_{600}$) from three time points (35-, 40- and 45-mins post induction). Values for the limit of detection (LOD) and Environmental Protection Agency (EPA) guideline values are indicated. Values are presented as mean and ± standard deviation of either two or three independent replicates.

-35 -10

$P_{merR20}$ ATGTATGAC**TTGACC**GTGTACTATGGTACAGGGTT**TATACT**TTTTATT

$P_{merR19}$ ATGTATGAC**TTGACC**GTGTACTATGGTACAGGGT**TATACT**TTTTATT

**Supplementary Figure S6. Wild-type ($P_{merR20}$) and mutant ($P_{merR19}$) promoters.** The wild-type $P_{merR20}$ promoter was used as a template for targeted mutagenesis to remove 1 bp adjacent to the -10 element (indicated by red arrow), as done previously by Parkhill *et al* [7] to generate the mutant promoter $P_{merR19}$. The MerR dyad sequence is underlined with both the -35 and -10 element in bold.

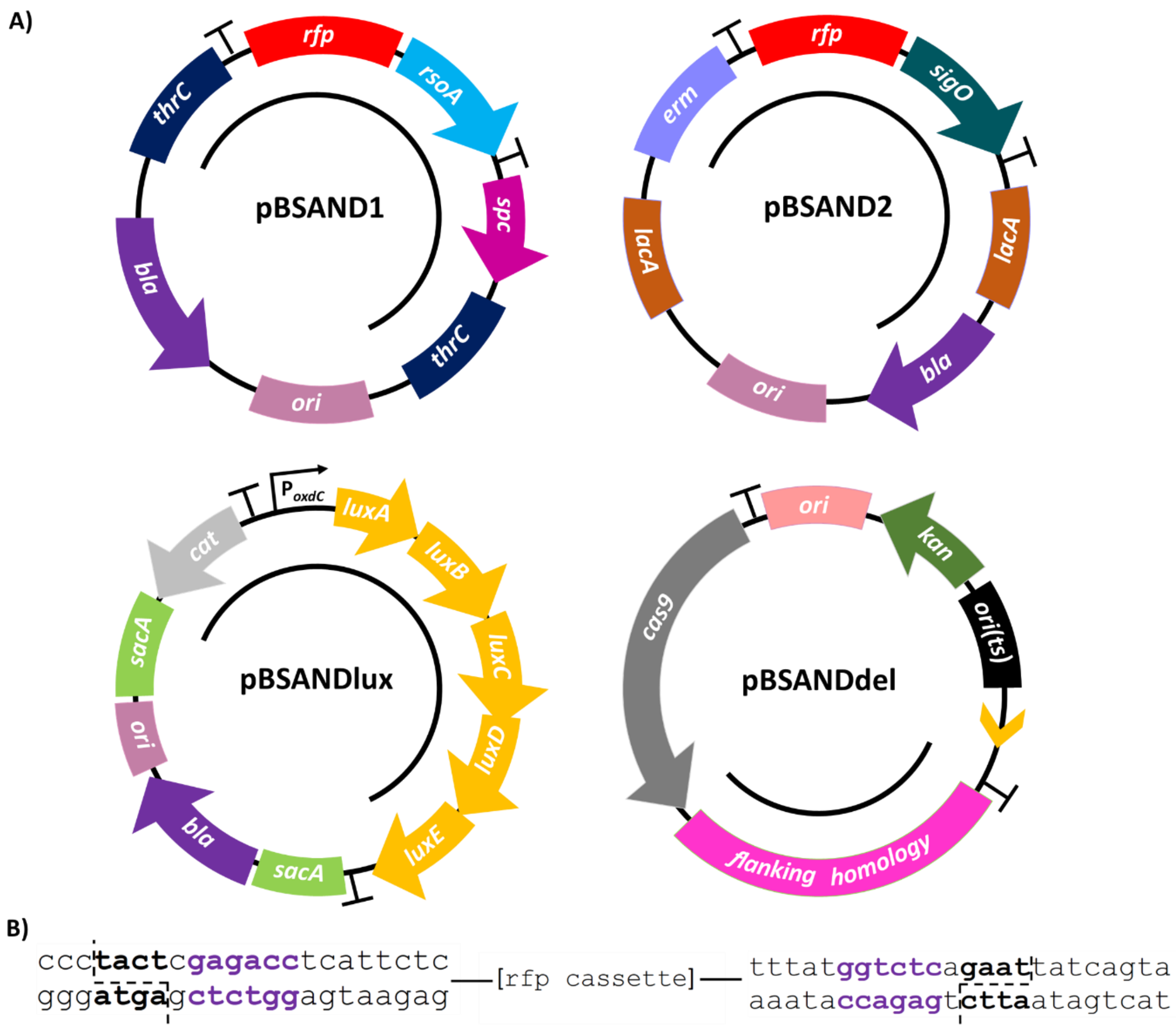


**Supplementary Figure S7. The *B. subtilis* SANDBOX plasmids. A)** Vector architecture for plasmids pBSAND1 and pBSAND2 both of which contain one half of the split sigma factor system SigO-RsoA. Plasmids pBSAND1, pBSAND2 and pBSANDlux are all integrative vectors with resistance markers *spc* (spectinomycin), *erm* (MLS; *macrolide, lincosamide and streptogramin B antibiotics if induced by erythromycin)* and cat (chloramphenicol) and integrate at the loci *thrC, lacA* and *sacA*, respectively. Whilst pBSANDdel is an integrative vector, the flanking homology region (shown in pink) is the only integrative portion of the plasmid. The gRNA to cut within the *sigO-rsoA* regulon is indicated with an orange arrowhead. Plasmid pBSANDlux is a luciferase-based reporter vector ($P_{oxdC}$-*luxABCDE*) and pBSANDdel is a modified CRISPR-Cas9 vector designed to knockout the SigO-RsoA regulon. The integrative portion of all the logic gate plasmids are shown with a black line, terminators are indicated with the "T" symbol, and all comprise a *bla* (ampicillin) resistance marker to allow for selection in *E. coli* – the exception of which is pBSANDdel which has a *kan* (kanamycin) marker for selection in both *E. coli* and *B. subtilis*. Plasmids pBSAND1, pBSAND2, pBSANDlux and pBSANDdel are derived from pBS4S, pBS2E, pBS3Clux and pJOE8999 [3,4] . **B)** The Golden Gate cloning site based on BsaI. The RFP cassette is flanked by two Golden Gate restriction sites, highlighted in bold purple, with the overhang indicated in bold black.

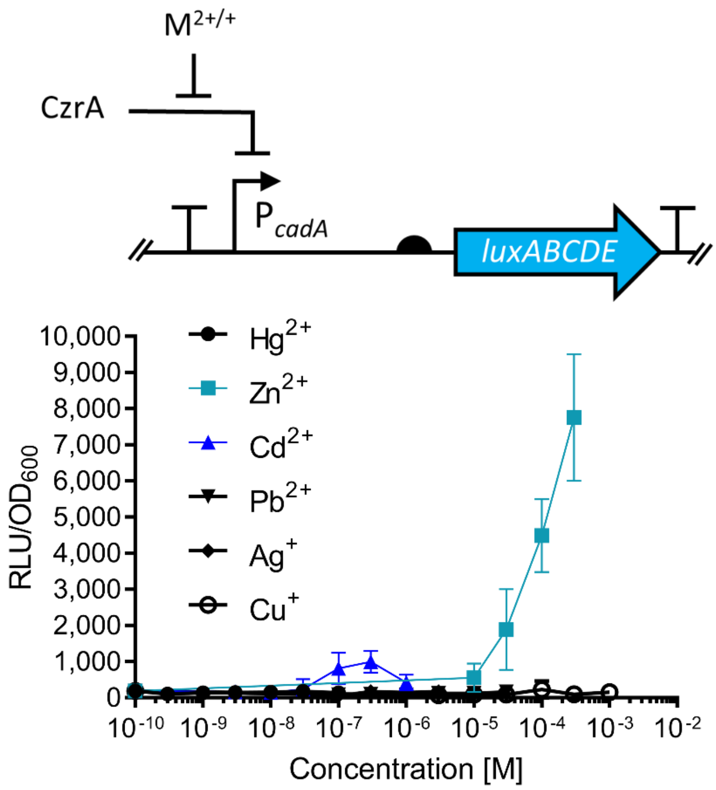


**Supplementary Figure S8. *Bacillus subtilis* natively regulated CzrA based circuit.** In the circuit shown, CzrA mediated repression of the cognate promoter $P_{cadA}$ is relieved upon the addition of heavy metal ions. Transcriptional output from $P_{cadA}$, measured via luciferase activity (luxABCDE, light blue arrow) is shown in response to various concentrations of heavy metals. Inducers, $Zn^{2+}$ and $Cd^{2+}$ are indicated in teal and dark blue respectively. $M^{+/2+}$ indicates the addition of either a monovalent or divalent metal ion. Cells were induced at $OD_{600}$ = ~ 0.03 with luciferase activity (relative luminescence units [RLU]) normalised to optical density ($OD_{600}$) values ($RLU/OD_{600}$) from three time points (35-, 40- and 45-mins post induction). Values are presented as mean and ± standard deviation of either two or three independent replicates.
